# Serum metabolome profiling in patients with mild cognitive impairment reveals sex differences in lipid metabolism

**DOI:** 10.1101/2024.11.11.623108

**Authors:** Rocio Diaz Escarcega, Vijay Kumar M. J., Vasilia E. Kyriakopoulos, Guadalupe J. Ortiz, Aaron M. Gusdon, Huihui Fan, Pedram Peesh, Maria P. Blasco Conesa, Gabriela Delevati Colpo, Hilda W. Ahnstedt, Lucy Couture, Stella H. Kim, Miriam Hinojosa, Christine M. Farrell, Sean P. Marrelli, Akihiko Urayama, Bhanu P. Ganesh, Paul E. Schulz, Louise D. McCullough, Andrey S. Tsvetkov

## Abstract

Alzheimer’s disease (AD) affects more women than men. Although women live longer than men, it is not longevity alone, but other factors, including metabolic changes, that contribute to the higher risk of AD in women. Metabolic pathways have been implicated in AD progression, but studies to date examined targeted pathways, leaving many metabolites unmeasured. Sex is often a neglected biological variable, and most metabolomic studies were not designed to investigate sex differences in metabolomic profiles. Here, we performed untargeted metabolomic profiling of sera from male and female patients with mild cognitive impairment (MCI), a common precursor to AD, and matched controls. We discovered significant metabolic changes in individuals with MCI, and found several pathways that were strongly associated with sex. Peptide energy metabolism demonstrated sexual dimorphism. Lipid pathways exhibited the strongest differences between female and male MCI patients, including specific phosphatidylcholine lipids, lysophospholipids, long-chain fatty acids, and monoacylglycerols. 1-palmitoleoyl glycerol and 1-arachidonoyl glycerol were higher in female MCI subjects than in male MCI subjects with no differences between control males and females. Conversely, specific dicarboxylic fatty acids were lower in female MCI subjects than male MCI subjects. In cultured astrocytes, 1-arachidonoyl glycerol promoted phosphorylation of the transcriptional regulator sphingosine kinase 2, which was inhibited by the transient receptor potential vanilloid 1 receptor antagonists, as well as chromatin remodelling. Overall, we identified novel sex-specific metabolites in MCI patients that could serve as biomarkers of MCI in both sexes, help further define AD etiology, and reveal new potential prevention strategies for AD.

**Highlights:**

▪ Individuals with MCI experience significant metabolic changes.
▪ Lipid pathways demonstrated the strongest sexual dimorphism in MCI.
▪ 1-monoacylglycerols showed a robust sex association, being higher in MCI females.
▪ Sex-specific metabolites may be MCI biomarkers in each sex.

## 1. Introduction

Alzheimer’s disease (AD) is the most common age-associated neurodegenerative disorder, and as human life expectancy increases, the prevalence of AD will also increase dramatically in the coming decades^1^. Newly developed anti-amyloid immunotherapies have shown modest clinical benefits but also have significant risks, including cerebral edema and intracerebral hemorrhage^2,3^, especially in women^4,5^, and they do not cure AD. Thus, more treatment approaches are urgently needed. Moreover, clinically relevant biomarkers are critically needed to identify the disease earlier and track disease progression, especially before irreversible or intractable pathology is established. The development of new therapeutic agents that target beneficial and/or deleterious signaling pathways involved in AD must be identified^6^.

AD has been intensively investigated with genomic, transcriptomic, proteomic, and metabolomic techniques to understand the molecular mechanisms of disease and to identify biomarkers^6^. For example, genes associated with AD have been discovered, such as the *ε*4 haplotype of apolipoprotein E (APOE)^7^. Analyses of various cell types in AD reveal that both the transcriptome and proteome are important in disease onset, progression, and severity^6^. Since metabolic changes in the brain and the peripheral tissues have been reported in AD^8^, targeted and untargeted metabolomics have been used to identify biomarkers and to establish causation for specific mechanisms that contribute to AD^6,9,10^.

Female sex is a risk factor for AD^11^. Although women live longer than men, older age is not the only contributing factor to the higher prevalence of AD in women^11^. The chromosomal complement, hormones, lifestyle, depression, and abnormal sleep behaviors may also be risk factors^12,13^. Female metabolism is highly affected by menopause^14,15^. Metabolomic data and sex differences in biomarkers in AD are poorly investigated and the underlying mechanisms in sex-specific pathways are not understood^11^.

Mild cognitive impairment (MCI) is also a major risk factor for AD. There is considerable variation in the progression of MCI to dementia. The conversion rate from MCI to dementia in 12 months is 5–39%^16–23^. Amnestic MCI with memory impairment may have a higher tendency than non-amnestic MCI (i.e., MCI with no memory impairment) to progress to AD^24^. Metabolic changes affect the conversion rate^25–27^. Sex differences and their underlying mechanisms in MCI are unclear^28^. The effect of sex on the circulating metabolites could reveal causative mechanisms of disease pathogenesis and progression and potential prevention strategies for AD. Clinically measurable molecular targets could also be used to identify sex-specific biomarkers as the basis for subsequent diagnostics and therapies for both male and female patients with MCI.

In this study, we performed untargeted metabolomic analyses of serum samples collected from male and female individuals with MCI and from healthy male and female controls. Our findings demonstrate sex-dependent metabolomic differences in patients with MCI versus healthy controls. Notably, peptide metabolism and energy metabolism showed sex differences. Lipid metabolism exhibited robust dimorphism according to sex in MCI cohorts, including specific dicarboxylic fatty acids, phosphatidylcholine lipids, lysophospholipids, long-chain fatty acids, and monoacylglycerol metabolites. We discovered two 1-monoacylglycerols-1-palmitoleoyl glycerol and 1-arachidonoyl glycerol (1-AG) that exhibit dimorphism according to sex in MCI individuals but showed no such sex differences in controls. Since astrocytes are important regulators of lipid homeostasis in health and disease^29^, we tested if 1-AG had any effect on cultured astrocytes. In astrocytes cultured from aged male and female mouse brains (22 months old), 1-AG induced phosphorylation of the transcriptional regulator sphingosine kinase 2 (SphK2), which was prevented by antagonists of the transient receptor potential vanilloid 1 (TRPV1) channel, a target of 1-monoacylglycerols^30^. 1-AG also promoted chromatin remodeling by altering G-quadruplex landscapes both in male and female astrocytes. Overall, we discovered novel sex-specific differences in metabolites among MCI individuals that add to our understanding of sex-dependent mechanisms of early cognitive impairment and offer new clinically relevant biomarker candidates for MCI in both sexes.

## 2. Methods

### 2.1. Patient sample selection

Serum samples included 20 male and female MCI patients and 20 male and female controls, matched by age, ethnicity, and sex (Supplemental Table 1). Four female patients with MCI were in hormone replacement therapy. Informed consent was obtained from all subjects.

### 2.2. Cognitive testing

Cognitive screens were used to aid early detection, severity grading, and longitudinal tracking of cognitive impairment. If scores fell below a recommended cut-off, further evaluation was typically completed to assess for a neurocognitive disorder based on the DSM-5 criteria. Various cognitive screens have been validated for use. The Mini-Mental Status Examination (MMSE) and the Montreal Cognitive Assessment (MoCA) are widely used screens that were developed with the goal of minimizing linguistic and cultural impact on scores and are available in different languages. The MMSE and MoCA were administered to the patient group, and the Mini-MoCA was used to screen control subjects for possible MCI.

The standard version of the MoCA is a 30-item tool that consists of tasks assessing executive functioning, visuospatial skills, language, attention, short-term memory, and orientation. One point was added to the total score if an individual had completed ≤12 years of education. Its psychometric properties are reliable for detecting possible MCI secondary to various neurological and medical conditions. A cut-off score of ≤25 (normal scores range from 26 to 30) produced a sensitivity of 90% and specificity of 87% for MCI^31^. Due to procedural challenges of in-person cognitive screening during the COVID-19 pandemic, control subjects were administered the Mini-MoCA by telephone. This 15-item version of the MoCA excludes tasks that are visually mediated or require use of paper and pencil. It has a recommended cut-off score of ≤11 for possible MCI (normal scores range from 12 to 15). A study of similar cognitive screens (telephone MoCA) demonstrated sufficient psychometric properties for screening MCI, with sensitivity and specificity set at 72% and 59%, respectively^32^.

The MMSE is a 30-item tool that was originally developed for dementia severity grading^33^. It has a lower sensitivity for detecting MCI, which may be partly due to the absence of tasks that can assess other cognitive domains, such as executive functioning^34^. The MoCA has better sensitivity without significantly compromising specificity and also correlates well with standalone neuropsychological measures^35,36^. Due to the variable use of the MMSE and MoCA in clinical and research settings, studies have proposed conversion tables using the equipercentile equating method. For this study, the MMSE scores in our patient group were converted to the MoCA for consistency with patient subjects who were originally administered the standard MoCA^37,38^.

### 2.3. Sample accessioning and preparation

Serum samples were collected under the approved IRB protocols (“BioRepository of neurological diseases and sex-specific immune responses to SARS-CoV-2 and chronic neurologic symptoms”, HSC-MS-17-0452; “Neutrophil-DEspR+ in COVID”, HSC-MS-20-0780, and “UT Health Neurocognitive Disorders Center Biobank” and the IRB number was HSC-MS-19-0219) were inventoried, assigned with a unique identifier, and stored at -80°C. This identifier was used to track all sample handling, experiments, and results. Serum samples were prepared using the automated MicroLab STAR® system from Hamilton Company. Proteins were precipitated with methanol under vigorous shaking for 2 min and followed by centrifugation. The resulting extract was divided into five fractions: two for analysis by two separate reverse phase (RP)/ultrahigh-performance liquid chromatography-tandem mass spectroscopy (UPLC-MS/MS) methods with positive ion mode electrospray ionization (ESI), one for analysis by RP/UPLC-MS/MS with negative ion mode ESI, one for analysis by HILIC/UPLC-MS/MS with negative ion mode ESI, and one sample was reserved for backup.

### 2.4. UPLC-MS/MS

All methods used a Waters ACQUITY UPLC and a Thermo Scientific Q-Exactive high-resolution/accurate MS interfaced with a heated electrospray ionization (HESI-II) source and Orbitrap mass analyzer operated at 35,000 mass resolution. The sample extract was dried and then dissolved in solvents compatible to each of the four methods. Each solvent contained a series of standards to ensure injection and chromatographic consistency. One aliquot was analyzed using acidic positive ion conditions, chromatographically optimized for more hydrophilic compounds. A C18 column (Waters UPLC BEH C18-2.1×100 mm, 1.7 µm) was used, as well as water and methanol, containing 0.05% perfluoropentanoic acid (PFPA) and 0.1% formic acid (FA). Another aliquot was analyzed using acidic positive ion conditions; however, it was chromatographically optimized for more hydrophobic compounds. The extract was eluted from the same C18 column using methanol, acetonitrile, water, 0.05% PFPA and 0.01% FA. Another aliquot was analyzed using basic negative ion optimized conditions with a separate dedicated C18 column. The extracts were eluted from the column using methanol and water, but with 6.5 mM ammonium bicarbonate, pH 8. The fourth aliquot was analyzed via negative ionization, after elution from an HILIC column (Waters UPLC BEH Amide 2.1×150 mm, 1.7 µm) using a gradient of water and acetonitrile with 10 mM ammonium formate, pH 10.8. The MS analysis alternated between MS and data-dependent MSn scans using dynamic exclusion.

### 2.5. Data extraction and compound identification standards

Raw data were extracted, peak-identified, and processed using Metabolon’s hardware and software. Compounds were identified by comparison to library data of purified standards or recurrent unknown entities. The library contains the retention time/index (RI), mass to charge ratio (m/z), and chromatographic data (including MS/MS spectral data) on all molecules in the library. More than 3300 commercially available purified standard compounds were acquired and registered. Additional mass spectral data were created for structurally unnamed biochemicals, which were identified by their chromatographic and mass spectral signatures.

A biochemical name with no symbol indicates a compound confirmed based on an authentic chemical standard, and Metabolon is highly confident in its identity (Metabolomics Standards Initiative Tier 1 identification). A biochemical name with an * indicates a compound that has not been confirmed based on a standard, but Metabolon is confident in its identity. A biochemical name with an ** indicates a compound for which a standard is not available, but Metabolon is reasonably confident in its identity. A biochemical name with a (#) or [#] indicates a compound that is a structural isomer of another compound in the Metabolon spectral library. For example, a steroid that may be sulfated at one of several positions indistinguishable by the mass spectrometry data or a diacylglycerol for which more than one stereospecific molecule exists. The term “biochemicals” refers to a molecular feature in the text. Partially metabolites are considered “Tier 3” or “Level 3” as defined by the Metabolomics Standards Initiative. A Tier 3 metabolite is identified based on its spectral characteristics and likely belongs to a specific chemical class without a definitive match to a known authentic standard.

### 2.6. Statistical methods

Standard statistical analyses were performed in Jupyter Notebook or R or JMP. Association analyses between metabolites and APOE genotypes were carried out using the non-parametric Wilcoxon signed-rank test in R software (Version 4.2.2).

### 2.7. Random forest

This supervised classification technique is based on an ensemble of decision trees^39^. For a given decision tree, a random subset of the data with identifying true class information was selected to build the tree (“training set”), and then, the remaining data, the “out-of-bag” variables, were passed down the tree to obtain a class prediction for each sample. This process was repeated thousands of times to produce the forest. The final classification of each sample was determined by computing the class prediction frequency (“votes”) for the out of bag variables over the whole forest. The prediction accuracy was an unbiased estimate of how well one can predict sample class in a new data set. R was used for the random forest method.

### 2.8. Principal components analysis

Principal components analysis (PCA) was an unsupervised analysis that reduced the dimension of the data. Each principal component was a linear combination of every metabolite, and the principal components were uncorrelated. The number of principal components was equal to the number of observations. The first principal component was computed by determining the coefficients of the metabolites that maximize the variance of the linear combination. The second component found the coefficients that maximize the variance with the condition that the second component was orthogonal to the first. The third component was orthogonal to the first two components, and so on. The total variance was defined as the sum of the variances of the predicted values of each component (the variance is the square of the standard deviation), and for each component, the proportion of the total variance was computed.

### 2.9. Linear regression analysis

A generalized linear model was fitted with a log-scaled normalized signal for each of the 1136 metabolites using R software to predict the MMSE and MOCA scores of MCI patients. P values < 0.05 were counted as significant. Correlations between 1-monoacylglycerol metabolites and MoCA/MMSE scores were visualized using ggplot2 R package.

### 2.10. Bioinformatics

All bioinformatics analyses were performed in R (R Foundation for Statistical Computing). Fold-changes (FC) were calculated for each metabolite, comparing control to MCI subjects (males and females). Changes in metabolites were considered significantly increased at FC > 2 and decreased at FC < 2 with false discovered rate (FDR)-corrected *P*-values < 0.05. Sparse partial least squared discriminant analysis (PLS-DA) was used with the mixOmics library in R (http://mixomics.org). Adding discriminant analysis to the PLS algorithm allowed for classification of large datasets^40,41^. We used this method to select the most discriminative metabolites to classify groups. The *perf* function was used to determine the number of components. The number of components to use to maintain a classification error rate less than 0.1 was determined using the *perf* function. The *tune.splsda* function was used to determine the number of metabolites to minimize the balanced error rate.

### 2.11. Chemicals and antibodies

Antibodies against SphK2 were from ECM Biosciences (#SP4621). Antibodies against phospho-SphK2 (phospho-Thr614) were from Aviva Systems Biology (#OASG06779). Antibodies against S1OOβ were from Abcam (#AB52642). Antibodies against AQP4 were from Sigma-Aldrich (#MABN2526). Antibodies against GFAP were from Novus Biologicals (#NBP1-05198). Secondary anti-rabbit Alexa Fluor 488-labeled (#A11008) and anti-chicken Alexa Fluor 546-labeled (#A11040) antibodies were from Life Technologies. The G4P sequence was published by the Tan lab^42^ (see acknowledgements), and the G4-mScarlet construct was published by us^43^. 1-AG (#35474-99-8), JNJ17203212 (#30930), and CAY10568 were from Cayman Chemicals (#10012565).

### 2.12. Primary culture of male and female astrocytes

Astrocytic cultures were prepared from 22-month-old male and female C57BL/6J mice. We used astrocytes from aged mice because our metabolomic analyses were performed on aged individuals. The brains were dissected, and the cerebellum was removed. Brain samples were enzymatically digested with the adult brain dissociation kit, according to the manufacturer protocol (MACS Miltenyi Biotec, #130-107-677) with minor modifications. Briefly, brain samples were digested at 37 °C for 20 min with gentle mixing every 10 min. Cells were then passed through a 70-µm-pore mesh, and debris was separated by centrifugation at 500 X g for 10 min. Single-cell suspensions were then plated onto a poly-D-lysine-coated 75-cm^2^ flasks (Sigma-Aldrich, #P6407) and cultured in Dulbecco’s modified Eagle’s medium/Ham’s F-12 1:1 (Cytiva; #SH30261.01) supplemented with 10% fetal bovine serum, 50 µg/mL streptomycin, and 50 U/mL penicillin (Gibco; #15140122). Astrocytes were cultured in 37 °C humidified incubators with 5% CO_2_. The medium was changed every 3 days.

For treatments, cells were cultured *in vitro* up to two passages. Astrocytes were then seeded onto multi-well glass slides coated with Matrigel and treated with vehicle (ethanol: PBS – 1:2), 4 µM 1-AG, and 1-AG in combination with TRPV1 inhibitors – 2 µM CAY10568, and 2 µM JNJ JNJ17203212 overnight. After treatment, astrocytes were fixed with 4% PFA, blocked with blocking buffer (5% normal goat serum), 1% BSA and 0.3% Triton-X-100 for 1 hr. After blocking, cells were treated with primary antibodies against SphK2 and phospho-SphK2, and GFAP overnight. The next day, cells were washed with 1X PBS three times and treated with secondary antibodies, mounted, and imaged using a confocal microscope (Nikon A1R confocal microscope).

For the experiments with the G4P-mScarlet reporter, male and female astrocytes were plated into a 6-well tissue-culture plate. At about 80% confluence, cells were infected with the G4P-mScarlet lentivirus (>10^8^ TU/ml). Astrocytes were imaged with Incucyte-ZOOM live-cells microscopy incubator (Essen Bioscience). Images were then analyzed with ImageJ.

## 3. Results

### 3.1. Selection of control and MCI patient samples

We analyzed serum samples from 20 male and 20 female control individuals and 20 male and 20 female patients with MCI. MCI was determined by clinical, cognitive, and functional criteria (see Methods). Patients with MCI were matched to controls based on age, race, and sex (Supplemental Table 1 and Methods).

### 3.2. Global metabolomic profiling showed differences between controls and MCI patients and between males and females

Control and MCI serum samples were loaded equivalently across the analytical platforms. In the serum sample dataset, 899 biochemicals were detected by ultrahigh-performance liquid chromatography-tandem mass spectroscopy. We then used PCA to detect similarities, differences, and patterns within the datasets. We discovered that one male control sample clearly separated from all other samples (Supplemental Figure 1a, b). Heatmaps were then generated with and without the separated male control sample (Supplemental Data 1). As evident in Supplemental Data 1, the removal of the separated sample did not affect the overall results.

ANOVA contrasts revealed that 179 biochemicals differed between control males and females, and 186 biochemicals differed between MCI male and MCI female patients. Furthermore, 178 biochemicals differed between control and MCI females, and 176 biochemicals differed between control and MCI males (Figure 1a). 89 biochemicals approached significance (0.05 < p < 0.10) between control males and females, and 65 biochemicals approached significance (0.05 < p < 0.10) between MCI male and MCI female patients. Between control and MCI females, 77 biochemicals approached significance (0.05 < p < 0.10), and between control and MCI males, 67 biochemicals approached significance (0.05 < p < 0.10) (Figure 1a and Supplemental Figure 2a, b). We grouped metabolites according to their super pathways. Metabolites fell into several metabolic pathways, including amino acids, carbohydrate and energy metabolites, peptides, lipids, nucleotides, cofactors and vitamins, as well as xenobiotics and partially characterized molecules (Figure 1b). Thus, multiple metabolites exhibited an interaction with sex and MCI status.

**Fig. 1.**
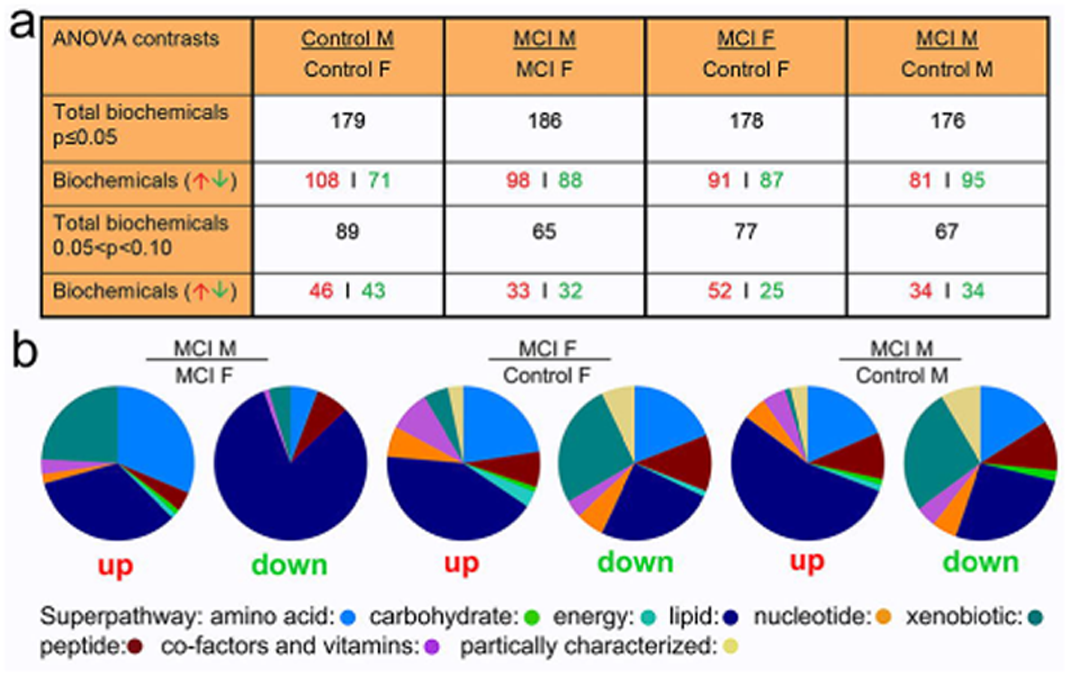
Metabolite summary and significantly altered biochemicals. **(a)** The dataset comprises 6,468 biochemicals. ANOVA contrasts were used to identify biochemicals that differed significantly between experimental groups. A summary of the numbers of biochemicals that achieved statistical significance (p≤0.05), as well as those approaching significance (0.05<p<0.10), is shown. P values were adjusted for multiple comparisons. Analysis by two-way ANOVA identified biochemicals exhibiting significant interaction and main effects for experimental parameters of MCI and sex. **(b)** A summary of biochemical families that achieved statistical significance (p≤0.05). Ten families of metabolic biochemicals were identified: amino acids, peptides, carbohydrates, energy, lipids including primary and secondary bile acid metabolites, nucleotides, co-factors and vitamins, xenobiotics, and partially characterized biochemicals. Partially metabolites are considered “Tier 3” or “Level 3” as defined by the Metabolomics Standards Initiative. A Tier 3 metabolite is identified based on its spectral characteristics and likely belonging to a specific chemical class but without a definitive match to a known authentic standard. See Methods for details.

### 3.3. Random forest analysis of metabolites

In machine learning, random forest analysis bins individual samples into groups based on metabolomic similarities and variances to show metabolomic biomarkers of interest^44^. Biochemical profiles from these random forest analyses predicted the controls versus MCI group (with male and female samples combined) with 100% accuracy (Figure 2a). Biochemical profiles predicted the male controls versus male MCI samples with an accuracy of 100% (Figure 2b). Finally, biochemical profiles predicted the female controls versus female MCI samples with an accuracy of 95% (Figure 2c).

**Fig. 2.**
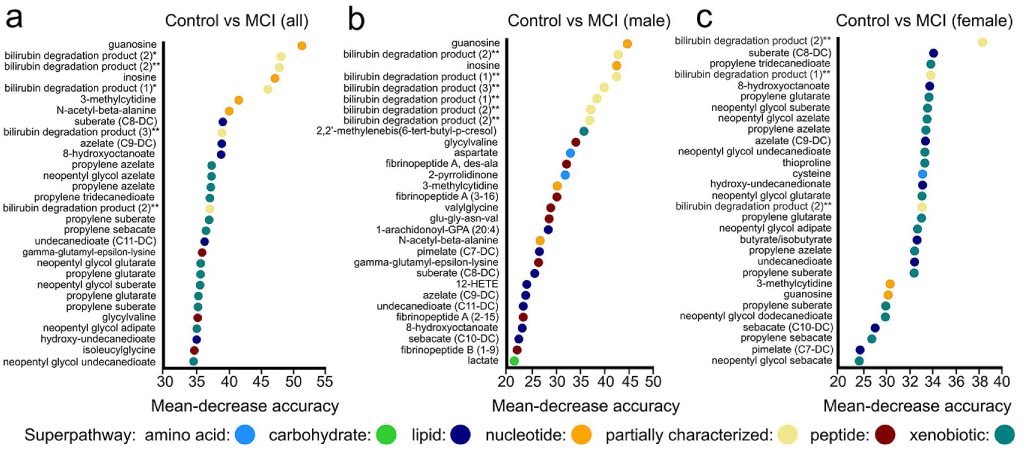
Random forest analysis of metabolic control versus MCI datasets. Random forest analysis bins individual samples into groups, based on their metabolite similarities and variances. **(a)** The biochemical profiles predict the control sample group versus MCI sample group with a predictive accuracy of 100%. Random forest analysis shows the 30 most-important metabolites. **(b)** The biochemical profiles predict the control male sample group versus male MCI sample group with a predictive accuracy of 100%. Random forest analysis demonstrates 30 top-ranking biochemicals of importance, based on control male and MCI male group separation in serum samples. **(c)** The biochemical profiles predict the control female sample group versus MCI female sample group with a predictive accuracy of 95%. Random forest analysis demonstrates the 30 top-ranking biochemicals of importance, based on control female and MCI female group separation in serum samples. A biochemical name with an * indicates a compound that has not been confirmed based on a standard, but we are confident in its identity. A biochemical name with an ** indicates a compound for which a standard is not available, but we are reasonably confident in its identity.

In the control versus MCI group (with male and female samples combined), the 30 top-ranking biochemicals were xenobiotics (13), lipid metabolism (5), partially characterized (5), nucleotide biochemicals (4), and peptides (3) (Figure 2a). In the male controls versus male MCI samples, the 30 top-ranking biochemicals were lipid metabolism (8), peptides (8), partially characterized (6), nucleotides (4), amino acid metabolites (2), a carbohydrate metabolite (1), and a xenobiotic metabolite (1) (Figure 2b). In the female controls versus female MCI samples, the 30 top-ranking biochemicals were xenobiotics (16), lipid metabolism (8), partially characterized (3), nucleotides (2), and an amino acid metabolite (1) (Figure 2c). Intriguingly, random forest analyses revealed multiple xenobiotics that fell within the top 30 metabolites in the female (control female-versus-MCI female cohorts) and all sample comparisons (control versus MCI (all)) (Figure 2a, c). In addition to xenobiotics, the major changes between the control and MCI groups were in lipids and their metabolites.

Next, we analyzed male-versus-female samples within the control and MCI groups. Biochemical profiles for the male control samples versus female control samples were predicted with 90% accuracy (Figure 3a). Biochemical profiles for the male MCI samples versus female MCI samples were predicted with 82.5% accuracy (Figure 3b). In the male control versus female control samples, the 30 top-ranking biochemicals were lipid metabolism (22), amino acid metabolism (4), cofactors and vitamins (2), a partially characterized molecule (1), and a xenobiotic metabolite (1) (Figure 3a). In the male MCI samples versus female MCI samples, the 30 top-ranking biochemicals were lipid metabolism (16), amino acid metabolism (10), peptide (2), an energy metabolite (phosphate) (1), and a nucleotide metabolite (1) (Figure 3b). The major differences between male and female samples in the control group, as well as between male and female samples in the MCI group, were lipids and their metabolites.

**Fig. 3.**
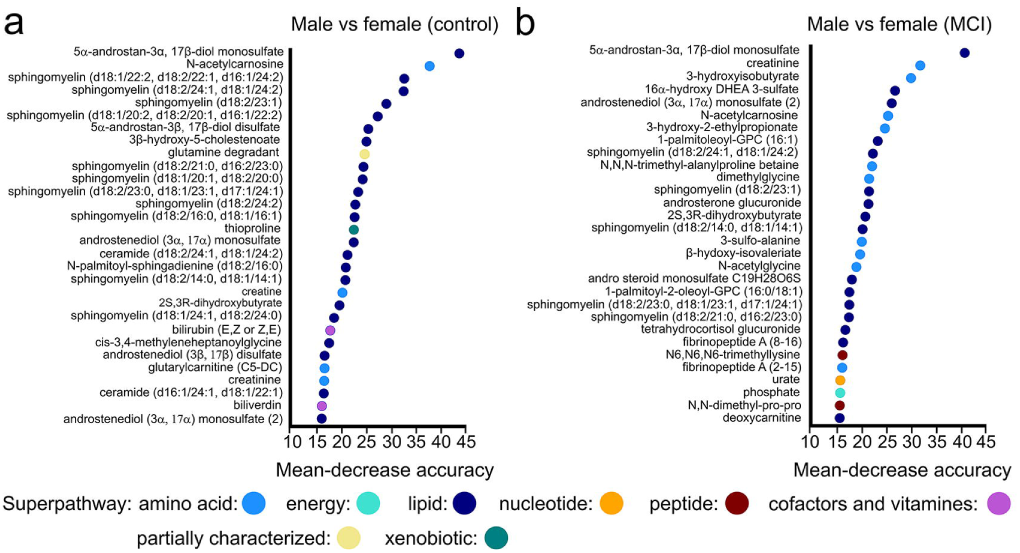
Random forest analysis of metabolic male versus female datasets. **(a)** The biochemical profiles are successful in predicting sex groups correctly with 90% accuracy (control male subjects versus control female individuals). Random forest analysis demonstrates the 30 most important metabolites in the control group (control male and female subjects). **(b)** The biochemical profiles predict the MCI male versus female sample group with a predictive accuracy of 82.5%. Random forest analysis demonstrates 30 top-ranking biochemicals of importance based on MCI male versus female group separation in serum samples.

### 3.4. Partial least squares-discriminant analysis

PLS-DA is a technique frequently used in metabolomics to classify metabolites^45^. Here, we used PLS-DA to determine the main metabolites contributing to differences between groups (Figure 4). We compared controls versus MCI patients (Figure 4a), control males versus MCI males (Figure 4b), and control females versus MCI females (Figure 4c) with the top 10 metabolites distinguishing the groups displayed in loading plots (Figure 4). The nucleosides guanosine and inosine were higher in the male MCI group than in the control male group, but the bilirubin degradation products and dipeptides (glycylvaline and valyglycine) were lower. Conversely, fatty acids (azelate, hydroxyundecanedioate, undecanedioate, 8-hydroxyoctanoate, pimelate, suberate) and xenobiotics (neopentylglycol, propylene azelate, propylene glutarate, propylene suberate) were lower in the female MCI group than the male MCI cohort.

**Fig. 4.**
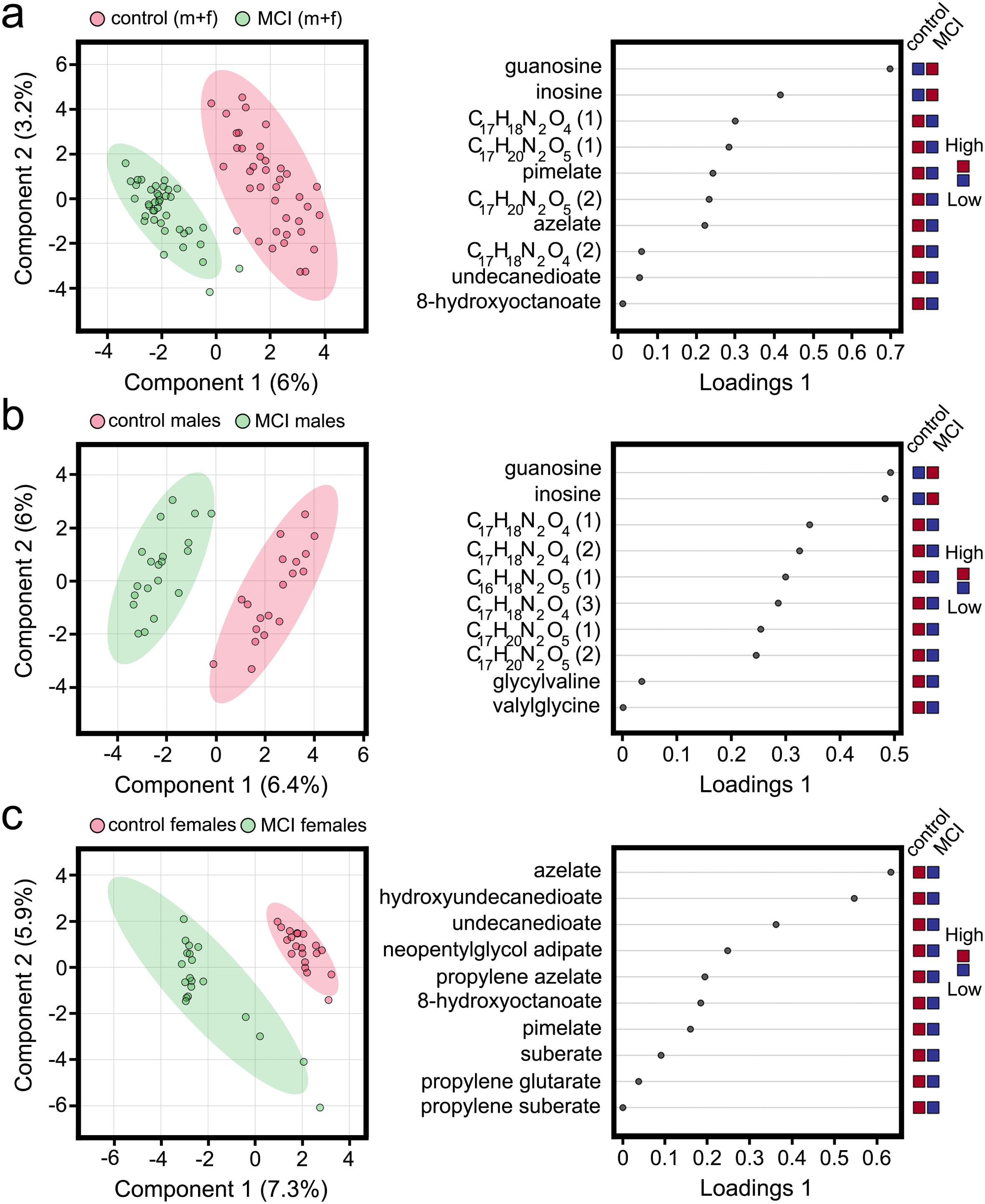
Partial least squares-discriminant analysis (PLS-DA) to determine the main metabolites contributing to differences between groups. **(a)** PLS-DA compared controls versus MCI patients with the top 10 metabolites distinguishing the groups displayed in loading plots on the right panel. **(b)** PLS-DA was performed to compare control males versus MCI males with the top 10 metabolites, distinguishing the groups displayed in loading plots on the right panel. **(c)** PLS-DA was performed to compare control females versus MCI females with the top 10 metabolites, distinguishing the groups displayed in loading plots on the right panel. A biochemical name with an ** indicates (bilirubin degradation products) a compound for which a standard is unavailable, but we are reasonably confident in its identity.

### 3.5. Several peptide metabolites show an association with sex and MCI status

A reduction in the extracellular levels of certain peptides (e.g., valylglycine) is associated with cell senescence^46^. In our data, several peptide metabolites were associated with sex and MCI status. In particular, dipeptides (e.g., leucylalanine and valylglycine) were lower in female and male MCI patients than same sex controls (Figure 5, Supplemental Data 1). Both peptides are lower in males with MCI than in females with MCI.

**Fig. 5.**
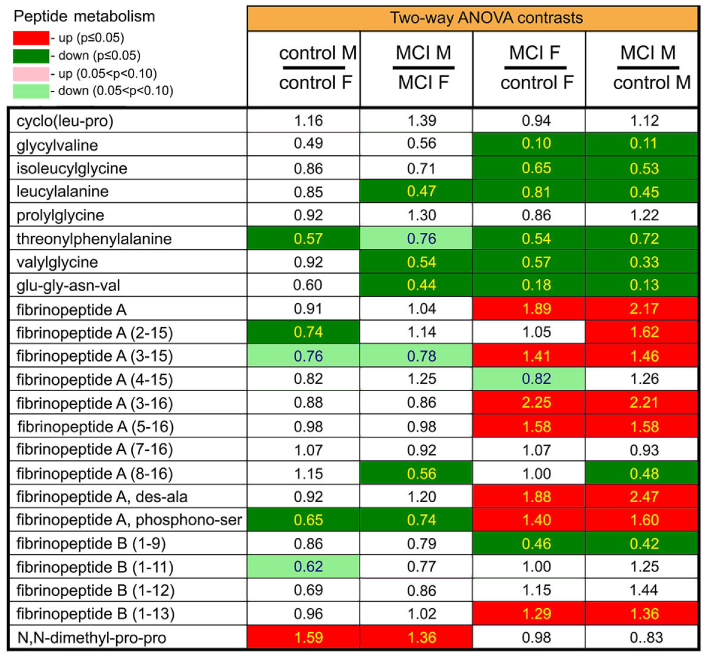
Several peptide metabolites show an association with sex and MCI status. Red and green cells indicate p≤0.05 (red indicates the fold-change values are significantly higher for that comparison; green values are significantly lower). Light red and light green shaded cells indicate 0.05<p<0.10 (light red indicates the fold-change values trend higher for that comparison; light green values trend lower). Note that there is a sex association for many metabolites.

Fibrinopeptides regulate coagulation^47^. We discovered that fibrinopeptide A cleavage fragments are higher in MCI female and male patients than in same sex controls (Figure 5, Supplemental Data 1). Levels of fibrinopeptide A increase with aging^47^, and we found that levels of fibrinopeptide A cleavage fragments were even higher in patients with MCI.

### 3.6. Energy metabolites succinate and phosphate exhibit an association with sex and MCI status

Succinate, fumarate, and malate are intermediates of the energy pathways in cells. We found that succinate levels were higher in male and female MCI patients than control male and female individuals. Succinate levels were higher in female MCI patients than male MCI patients (Supplemental Figure 3, Supplemental Data 1). Levels of fumarate and malate were also higher in female MCI patients than controls with no changes in the male cohorts. Finally, phosphate levels were lower in females with MCI than control females. MCI females had lower levels of phosphate than male MCI patients (Supplemental Figure 3, Supplemental Data 1). Our data suggest that energy pathways contribute to sex differences in MCI.

### 3.7. Several middle-chain dicarboxylate fatty acids show an association with sex and MCI status

Dicarboxylate fatty acids are metabolites with two carboxyl groups generated by the oxidation of fatty acids. They have a role in mitochondrial function and oxidative stress^48,49^. Out of 30 detected dicarboxylate fatty acids, only one exhibited a significant difference between control males and control females: dodecadienoic acid was lower in males (Figure 6, Supplemental Data 1). Pimelic, suberic, azelaic, sebacic, and undecanedioic acids were considerably lower in MCI females than control females and lower in MCI males than control males. Longer-chain dicarboxylate fatty acids (C12–C18) were higher in MCI females and MCI males than control females and males (Figure 6, Supplemental Data 1).

**Fig. 6.**
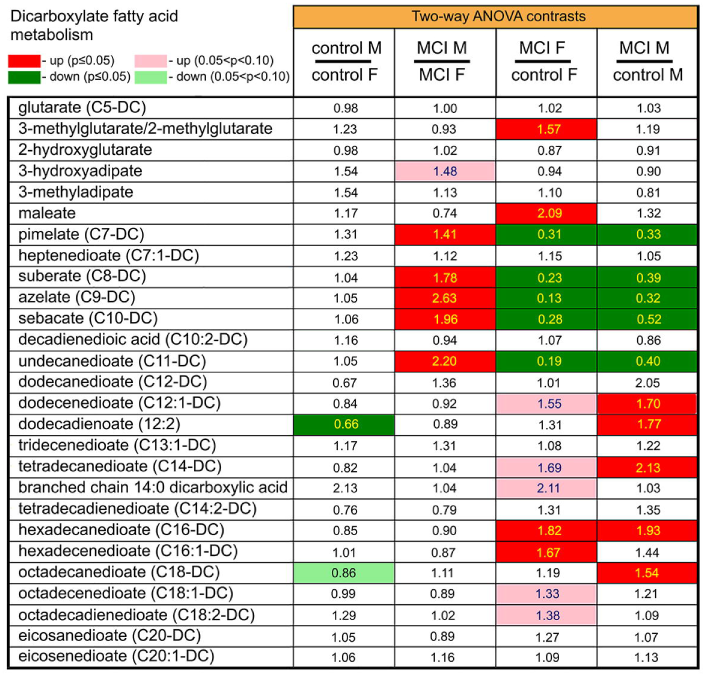
Differences in metabolism of dicarboxylate fatty acids between control subjects and MCI patients. Red and green cells indicate p≤0.05 (red indicates the fold-change values are significantly higher for that comparison; green values are significantly lower). Light red and light green shaded cells indicate 0.05<p<0.10 (light red indicates the fold-change values trend higher for that comparison; light green values trend lower). Note that there is a sex association for many metabolites.

Five dicarboxylate fatty acids were higher in MCI males than MCI females (pimelic, suberic, azelaic, sebacic, and undecanedioic acids) (Figure 6, Supplemental Data 1). Thus, middle-chain dicarboxylate fatty acids exhibited an association with sex in the MCI cohorts, which may mediate differences in oxidative pathways and mitochondrial function between males and females in MCI.

### 3.8. Phosphatidylcholine lipids exhibit a considerable association with sex and MCI status

Phosphatidylcholine (PC) lipids, a family of phospholipids that have choline as a headgroup, are membrane lipids. PC lipid types and levels change during aging with species and tissue specificity^50^. Two PC lipids achieved significance and were lower in the control males than control females (1-myristoyl-2-arachidonoyl-GPC (14:0/20:4) and 1-palmitoyl-2-docosahexaenoyl-GPC (16:0/22:6)) (Figure 7, Supplemental Data 1). MCI females had higher levels of PC lipids than control females, but this effect was less pronounced in the male cohort (p<0.05).

**Fig. 7.**
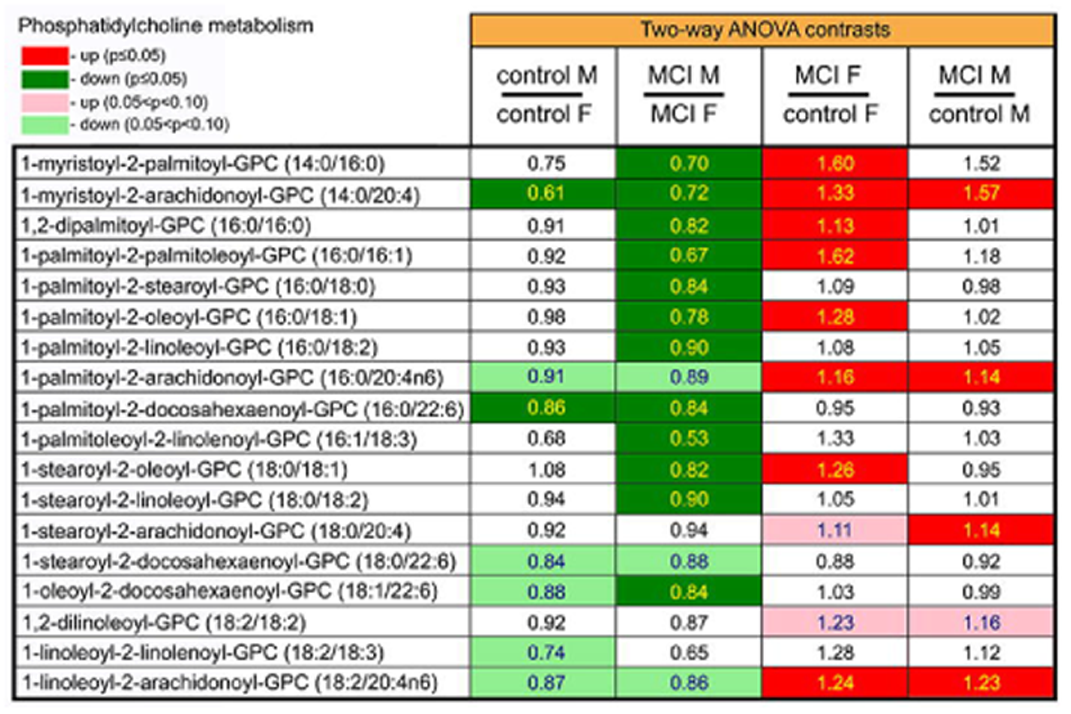
Differences in phosphatidylcholine metabolites between control subjects and MCI patients. Red and green cells indicate p≤0.05 (red indicates the fold-change values are significantly higher for that comparison; green values are significantly lower). Light red and light green shaded cells indicate 0.05<p<0.10 (light red indicates the fold-change values trend higher for that comparison; light green values trend lower). Note that phosphatidylcholine metabolites are lower in male than female MCI samples. Also, note that phosphatidylcholine metabolites are higher in female MCI patients than in controls.

Levels of PC lipids exhibited a strong sex association in the MCI group. Two-thirds of detected PC lipids were lower in male than the female MCI patients (Figure 7, Supplemental Data 1). Overall, many PC lipids exhibited a strong association with sex in MCI individuals, and as these lipids have a role in metabolic diseases^51^, their sex-association in MCI patients may contribute to changes in metabolism between MCI males and females.

### 3.9. Lysophospholipids were different in male and female subjects with MCI

Lysophospholipids are bioactive lipids characterized by a single carbon chain and a polar head group. Circulating lysophospholipids have important roles in immunity and inflammation^52^. We discovered that four lysophospholipids were lower in the male than the female control groups (1-linoleoyl-GPA (18:2), 1-stearoyl-GPE (18:0), 1-stearoyl-GPE (18:0), 1-stearoyl-GPI (18:0)) (Figure 8, Supplemental Data 1). There were minor differences in lysophospholipids between both male and female control and MCI groups (Figure 8, Supplemental Data 1).

**Fig. 8.**
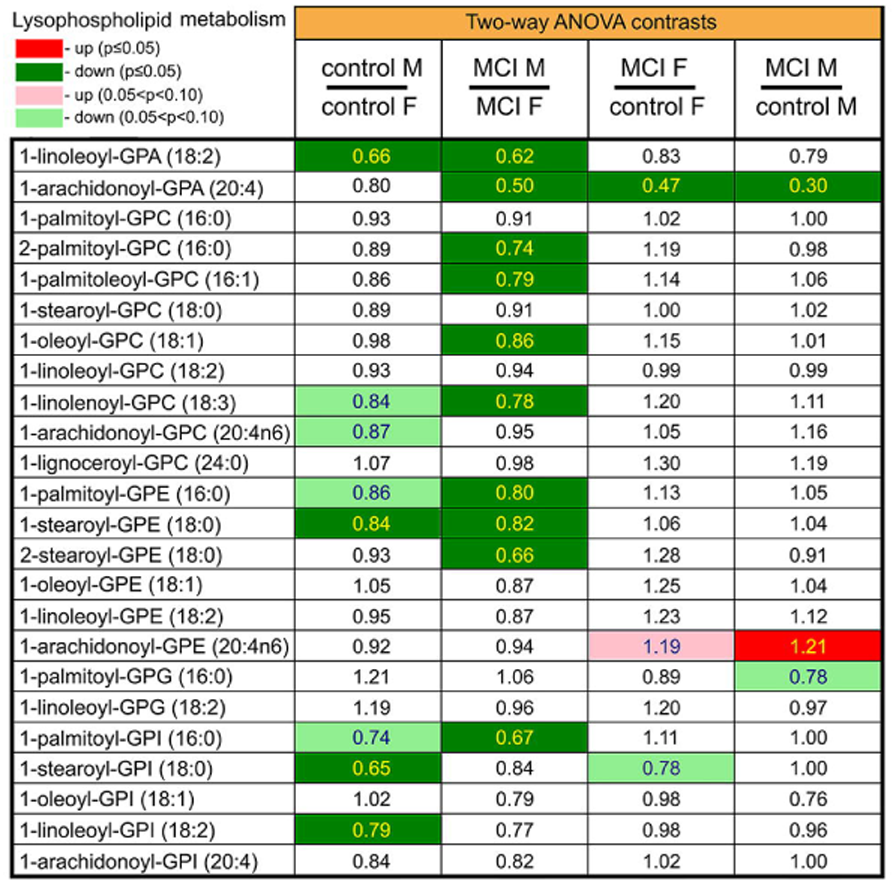
Lysophospholipids exhibit a strong association with sex and MCI status. Red and green cells indicate p≤0.05 (red indicates the fold-change values are significantly higher for that comparison; green values are significantly lower). Light red and light green shaded cells indicate 0.05<p<0.10 (light red indicates the fold-change values trend higher for that comparison; light green values trend lower). Note that many metabolites are considerably reduced in male MCI patients.

Ten lysophospholipids exhibited sex associations in MCI patients, with all being lower in males (Figure 8, Supplemental Data 1). Lysophospholipids exhibited a well-pronounced association with sex in the MCI cohort. Many lysophospholipids are pro-inflammatory molecules^52^, and their higher levels in MCI females may contribute to the increased levels of inflammation often observed in AD females, compared to AD males^53^.

### 3.10. Long-chain fatty acids exhibit differences between male and female subjects with MCI

Long-chain fatty acids contain 13–21 carbon atoms and have roles in diabetes, insulin resistance, and obesity, among others^54^. We detected 27 long-chain fatty acids in our samples. Stearidonic acid (18:4n3) and docosahexaenoic acid (DHA; 22:6n3) were significantly lower in control males than in control females (Figure 9, Supplemental Data 1). Four long-chain fatty acids were higher in MCI females than control females (palmitate (16:0), palmitoleate (16:1n7), tetradecadienoate (14:2), docosapentaenoate (n3 DPA; 22:5n3)) (Figure 9, Supplemental Data 1). Eight were significantly lower in MCI male than in MCI female patients. Overall, our data suggest that long-chain fatty acids are strongly associated with sex and MCI status, which may contribute to potential metabolic differences in MCI males and females.

**Fig. 9.**
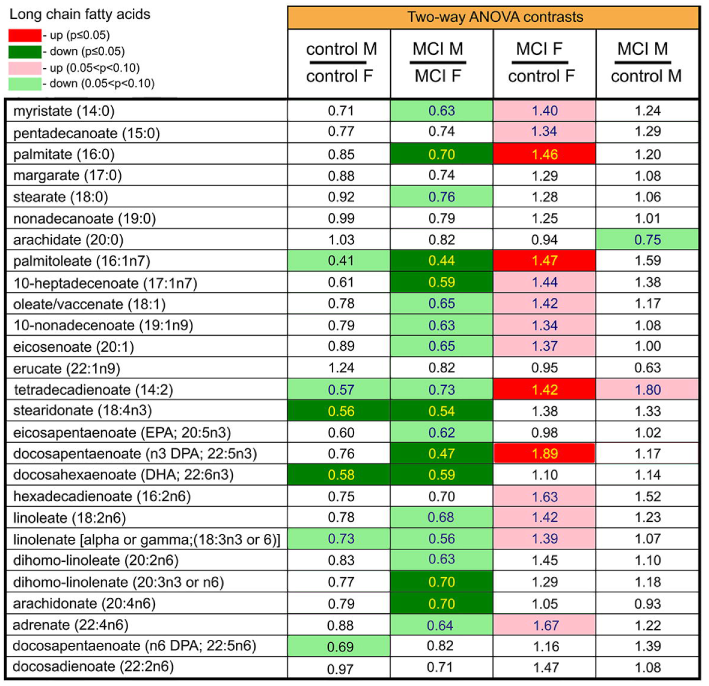
Differences in long-chain fatty acids between control subjects and MCI patients. Red and green cells indicate p≤0.05 (red indicates the fold-change values are significantly higher for that comparison; green values are significantly lower). Light red and light green shaded cells indicate 0.05<p<0.10 (light red indicates the fold-change values trend higher for that comparison; light green values trend lower). Note that there is a sex association for many metabolites.

### 3.11. 1-monoacylglycerols show a considerable association with sex and MCI status

Monoacylglycerols are a family of glycerides that consist of a molecule of glycerol linked to a fatty acid. They regulate immunity and metabolism^55,56^. We detected 12 1-monoacylglycerol metabolites, and none of them differed between control male and female individuals (Figure 10, Supplemental Data 1). Three metabolites (1-palmitoylglycerol (16:1), 1-linoleoylglycerol (18:2), 1-AG (20:4)) were higher in MCI female patients than in control females (Figure 10, Supplemental Data 1). Three metabolites (1-palmitoylglycerol (16:0), 2-oleoylglycerol (18:1), 2-linoleoylglycerol (18:2)) were lower in male MCI patients than in control male individuals (Figure 10, Supplemental Data 1). Intriguingly, 1−arachidonylglycerol (20:4) levels were higher in MCI females on hormone replacement therapy than in MCI females without hormone replacement therapy (p = 0.011, the Wilcoxon test).

**Fig. 10.**
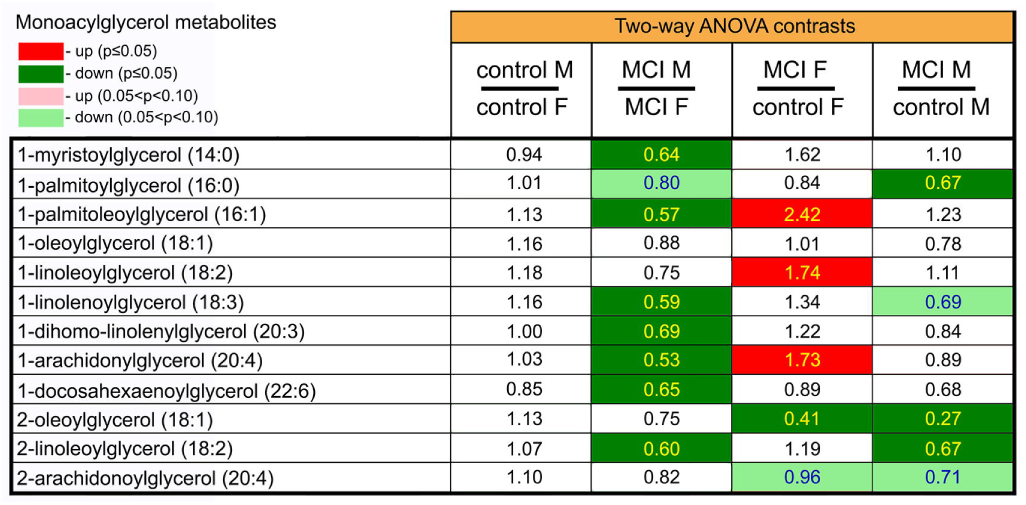
Monoacylglycerol metabolites differ between control subjects and MCI patients. Red and green cells indicate p≤0.05 (red indicates the fold-change values are significantly higher for that comparison; green values are significantly lower). Light red and light green shaded cells indicate 0.05<p<0.10 (light red indicates the fold-change values trend higher for that comparison; light green values trend lower). Note that there is a sex association for many metabolites.

About 2/3 of the detected 1-monoacylglycerol metabolites exhibited an association with MCI status and sex. MCI male patients had these metabolites at a lower level than MCI female patients. None of 1-monoacylglycerol metabolites differed between control male and female individuals, but there was a strong association of these metabolites to MCI status and sex, which can contribute to sex-associated differences in immune and metabolic function in MCI individuals.

### 3.12. MCI- and sex-associated lipid metabolites do not correlate with APOE status

We then determined if MCI- and sex-associated lipid metabolites correlated with APOE genotypes. Individuals with MCI were grouped by their sex and APOE genotypes (Supplemental Data 2), and groups with more than five individuals were retained for analyses. The non-parametric Wilcoxon signed-rank test was applied to assess the significance of APOE status for each lipid sex-specific metabolite in the MCI cohorts (Supplemental Data 3). Three lipid metabolites (suberate (C8-DC), sebacate (C10-DC), and 1-myristoylglycerol (14:0)) were significantly different between the APOE genotypes 4/4 and 3/4, whereas one metabolite (1-arachidonoyl-GPA (20:4)) differed between the genotypes 4/4 and 3/3 (p<0.05). However, no metabolite showed a significant difference after FDR corrections. Separate analyses of MCI females and males in the APOE cohorts showed no significant differences before or after FDR corrections (Supplemental Data 3). We, therefore, conclude that sex-dependent lipid metabolites in the MCI cohorts likely result from sex-associated lipid pathways rather from the APOE status of MCI individuals.

### 3.13. 1-AG enhances SphK2 phosphorylation and promotes chromatin remodeling in cultured male and female astrocytes

Unlike 2-monoacylglycerols^57^, 1-monoacylglycerol lipids are poorly studied, and their sex-associated functions are understood even less. Since 1-monoacylglycerols are considerably elevated in female MCI individuals (Figure 10), we hypothesized that these lipids could be pro-aging and pro-senescence metabolites, affecting cognition. We hypothesized that 1-AG may activate SphK2, a lipid kinase that modulates transcription, amyloid formation, and cellular senescence^58,59^. We used primary astrocytes cultured from aged male and female mouse brains (22-month-old mice) as an *in vitro* aging model. The purity of astrocytic cultures was confirmed with staining for astrocytic markers S100β and AQP4 (Supplemental Figure 4). Male and female astrocytes were treated with vehicle or 1-AG, fixed, and stained with antibodies against SphK2 and a phosphorylated form of SphK2 (phospho-Thr614). Since 1-monoacylglycerols can bind to and activate the TRPV1 channel^30^, we co-treated male and female astrocytes with 1-AG and TRPV1 antagonists, CAY10568 and JNJ-17203212 in parallel experiments. We discovered that 1-AG had marginal effects on SphK2 expression both in male and female astrocytes (Figure 11, Supplemental Figure 5). 1-AG induced SphK2 phosphorylation in the nucleus in male and female cells, and CAY10568 and JNJ-17203212 prevented SphK2 phosphorylation (Figure 11, Supplemental Figure 5). We then compared male versus female groups in the SphK2 and phospho-SphK2 groups and discovered that 1-AG had no effect on pan SphK2 levels, when compared to male and female cells, and exerted a much stronger effect on SphK2 phosphorylation in female astrocytes (e.g., higher phospho-SphK2 levels) than in male astrocytes, implicating sex differences at a molecular level in astrocytes cultured from aged mice (Figure 12).

**Fig. 11.**
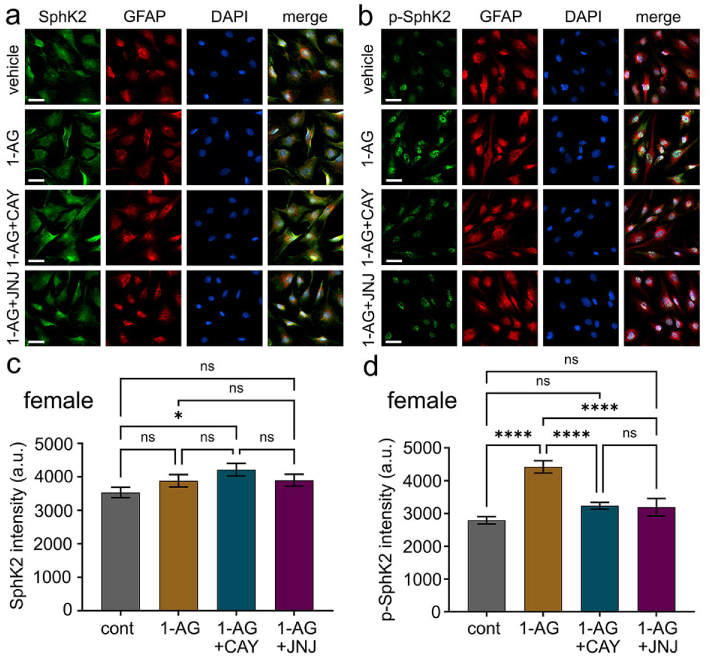
1-AG enhances SphK2 phosphorylation in aged mouse female astrocytes. **(a)** Female astrocytes were treated with control (vehicle), 1-AG alone (4 µM) or TRPV1 inhibitors (2 μM CAY10568 and 2 μM JNJ17203212), overnight. Astrocytes were fixed and stained with antibodies against SphK2 (green), GFAP (red), and with the nuclear DAPI dye (blue), and imaged. Bar, 10 µm. **(b)** Male astrocytes were treated with control (vehicle), 1-AG alone (4 µM) or TRPV1 inhibitors (2 μM CAY10568 and 2 μM JNJ17203212, overnight). Astrocytes were fixed and stained with antibodies against p-SphK2 (green), GFAP (red), and nuclear DAPI dye (blue), and imaged. Bar, 10 µm. **(c)** SphK2 intensities were measured from (a) with ImageJ (arbitrary units). *p=0.03, n.s., non-significant (ANOVA). At least 250 cells were analyzed from three experiments. **(d)** p-SphK2 intensities were measured from (b) with ImageJ (arbitrary units). ****p<0.0001, n.s., non-significant (ANOVA). At least 250 cells were analyzed from three experiments.

**Fig. 12.**
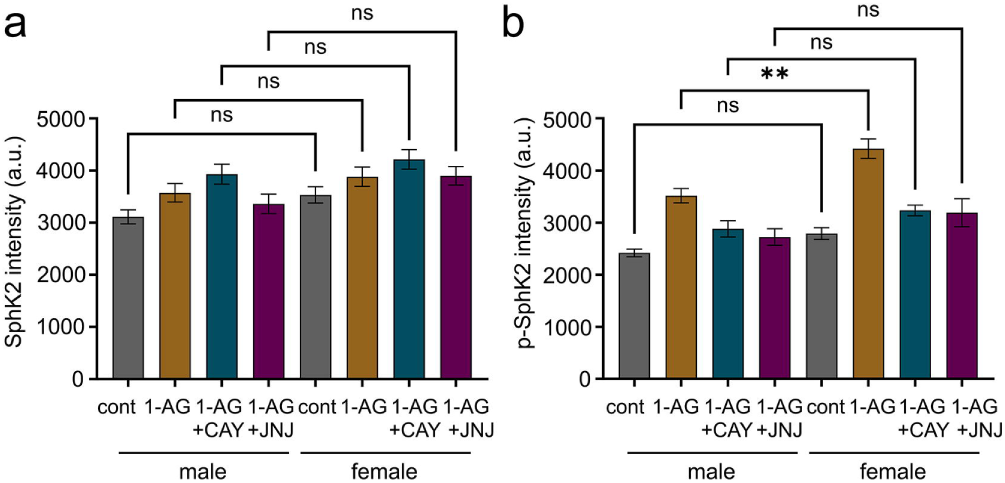
1-AG induces SphK2 phosphorylation in female astrocytes more potently than in male astrocytes. **(a)** 1-AG treatment (4 µM, overnight) does not differentially affect SphK2 in cultured male and female astrocytes. The data are from Figure 11 and Supplementary Figure 5. n.s., non-significant (one-way ANOVA). **(b)** 1-AG treatment (4 µM, overnight) induces SphK2 phosphorylation more potently in cultured female astrocytes than in male astrocytes. The data are from Figure 11 and Supplementary Figure 5. **p<0.01 (one-way ANOVA). More than 250 astrocytes per group were analyzed from three experiments.

Chromatin undergoes structural changes during aging, including alterations in G-quadruplex DNA landscapes^60,61^. We, therefore, hypothesized that 1-AG regulates G-quadruplex landscapes in male and female astrocytes cultured from age mice. Male and female astrocytes were infected with a lentivirus that encodes the fluorescent reporter of G-quadruplex DNA, the G4P-mScarlet^43^, and these cells were then treated with a vehicle or 1-AG and imaged with *in vivo* microscopy. 1-AG promoted changes in G-quadruplex landscapes (Supplemental Figure 6), and it did so more in male cells. We conclude that, in male and female astrocytes, the 1-monoacylglycerol 1-AG enhanced SphK2 phosphorylation and altered G-quadruplex DNA landscapes, likely affecting astrocytic transcription and replication in a sex-specific fashion.

### 3.14. Most sphingosines, dihydroceramides, ceramides, and sphingomyelins exhibit a strong association with sex, but not with MCI status

Sphingosines and ceramides are important lipids with a myriad of diverse functions^62^. Sphingosine and sphingosine-1-phosphate (S1P) were lower in MCI females compared to female controls (Supplemental Figure 7, Supplemental Data 1). S1P was lower in male MCI patients versus male controls. Sphinganine was lower in control males versus females and lower in MCI females versus controls. Sphinganine-1-phosphate was lower in MCI male patients versus females and was lower in MCI male patients versus controls (Supplemental Figure 7, Supplemental Data 1).

Most ceramides, including dihydroceramides, hexosylceramides, and lactosylceramides, exhibit a strong association with sex but not with MCI status. Their levels were lower in control and MCI males than in corresponding control and MCI females (Supplemental Figure 7, Supplemental Data 1). Notably, the levels of four ceramides were higher in males with MCI versus controls (ceramide (d18:1/14:0, d16:1/16:0), ceramide (d18:1/20:0, d16:1/22:0, d20:1/18:0), ceramide (d16:1/24:1, d18:1/22:1), glycosyl-N-stearoyl-sphingosine (d18:1/18:0); Supplemental Figure 7, Supplemental Data 1). Overall, with a few exceptions, most ceramides exhibited an association with sex but not with MCI status.

Sphingomyelins are abundant lipids that enter the circulation from the liver or intestine^63^. Their levels are regulated by sphingomyelinase secreted by vascular endothelial cells^64^. Most of the 29 sphingomyelins in our samples exhibited an association with sex. In particular, control males had lower levels of sphingomyelins than control females, and MCI males had lower levels of sphingomyelins than MCI females (Supplemental Figure 8, Supplemental Data 1). Male MCI patients had more of several sphingomyelins than controls (Supplemental Figure 8, Supplemental Data 1). Overall, sphingomyelins had a low association with MCI status and a strong dependency on sex: they were lower in control and MCI males than females.

### 3.15. MCI-associated metabolites correlate with MCI severity

Next, we examined the relationships between MCI- and sex-associated lipid metabolites and the MMSE or MoCA scores in MCI patients. We used a generalized linear model fitted with a log-scale-normalized signal for each of the metabolites using R software to predict the MMSE and MoCA scores of MCI patients (p<0.05). In the MCI cohort, we identified 40 metabolites associated with the measures of MMSE (Supplemental Data 4) and 37 that were associated with the MoCA scores (Supplemental Data 5). Four females with MCI on hormone replacement therapy did not cognitively differ from MCI females with no therapy (MMSE, p=0.27, the Wilcoxon test). Interestingly, three metabolites, including two 1-monoacylglycerols (1-myristoylglycerol (14:0) and 1-linolenoylglycerol (18:3)) correlated with both the MMSE and MoCA scores (Supplemental Data 4 and 5, Supplemental Figures 9 and 10).

## 4. Discussion

This study examined metabolomic profiles in serum samples of male and female MCI patients and controls. We found that peptide metabolism, including dipeptides and fibrinogen cleavage peptides, and energy pathways, were associated with sex in MCI patients. We also discovered strong dimorphism according to sex in lipid metabolic pathways, including dicarboxylic fatty acids, phosphatidylcholine lipids, lysophospholipids, long-chain fatty acids, and 1-monoacylglycerol metabolites. Individual lipid metabolites differed between male and female MCI patients, and whole pathways were often altered/shifted. Importantly, we identified two bioactive monoacylglycerols (1-palmitoleoyl glycerol and 1-AG) that were significantly higher in female MCI individuals than in male MCI individuals with no differences between control males and females. In astrocytes cultured from male and female 22-month-old mice, 1-AG promoted phosphorylation of SphK2, which was inhibited by the TRPV1 receptor antagonists, suggesting 1-AG signalling via the TRPV1 receptor, as well as chromatin remodelling. Overall, our studies provide evidence that MCI patients exhibit considerable metabolomic changes compared to controls, and several important sexually dimorphic pathways could serve as sex-specific biomarkers in MCI, a major risk factor for AD^16–23^.

Although the most established biomarkers for AD are β-amyloid and tau protein^65–68^, their predictive power for disease onset and progression is often poor, and their error in predicting the course of AD can be many years^69–71^. However, plasma pTau217 levels correlated well with concurrent AD pathophysiology in the brain and with prospective cognitive performance^72^. Plasma pTau217 and Aβ 42/40 levels predict the development of Aβ pathology in individuals with early stages of subthreshold Abeta accumulation^73^. A recent study showed that a commercially available pTau217 immunoassay accurately determined AD pathology and detected longitudinal changes, including ones at the preclinical stage^74^. Similarly, neurogranin protein, which has served as a CSF marker for synaptic loss, was proposed to be a biomarker for early symptomatic AD^75^, but has not shown utility as a biomarker for disease activity. In CSF, APOE, a fatty acid-binding protein, and the neuronal pentraxin receptor are associated with AD^76^. The D-serine/total serine ratio increases in the hippocampus of women with AD, and the insulin pathway is less active in women than men with AD^77^. A comprehensive study discovered 150 plasma and 154 CSF metabolites that were altered in AD patients, but 80% of the patients examined were men^78^, making those studies underpowered for examining sex differences. Plasma exosomal miRNAs may also serve as biomarkers in AD^79^, but their clinical utility has not been demonstrated. Apolipoproteins and their ratios have also been associated with the risk of developing dementia^80^. Since AD is a sexually dimorphic condition^81^, sex-specific biomarkers of AD are also urgently needed as sex-specific biomarkers may offer better predictive power than sex-nonspecific biomarkers.

Amyloid plaque burden does not correlate well with AD symptoms^82^; however, dysfunctional glucose metabolism and ATP biosynthesis (e.g., glycolysis, the tricarboxylic acid cycle (TCA), and the electron transport chain ATP synthesis) are defective and well correlated with the pathology, progression, and symptoms of AD^83^. In aging female rats, there is a metabolic transition from glucose utilization to catabolism of amino acids, fatty acids, lipids, and ketone bodies^84^. In our data, the levels of succinate are higher in male and female MCI patients than controls and higher in female than male MCI patients. Succinate is a metabolic intermediate of the electron transport chain, and its accumulation is linked to the defective TCA cycle and leads to epigenetic changes^85^ and inflammation^86^. Levels of fumarate and malate were also higher in female MCI patients than in control female individuals, which also indicates defects in the TCA cycle, epigenetic changes, and DNA damage^87,88^. Levels of phosphate are considerably lower in female MCI patients than in control female individuals, further indicating deficits in energy pathways. Female MCI patients have lower phosphate levels than males, demonstrating sex differences in energy metabolism. Thus, there are important sex differences in energy pathways in MCI patients.

Dicarboxylic acids, which are generated via the oxidative breakdown of fatty acids, regulate energy metabolism, mitochondrial function, and the immune system^48^. Excitingly, a recent paper demonstrated that the levels of pimelic (C7), suberic (C8), azelaic (C9), and sebacic (C10) acids are increased in urine samples from AD individuals. A lower hippocampal volume correlated with higher levels of C7–9 acids in urine^89^. The higher levels of azelaic acid in urine negatively correlated with Aβ42 and positively correlated with tau levels in CSF^89^. The authors concluded that a correlation between C7–9, AD biomarkers in CSF, and brain MRIs support their hypothesis that the excretion of C7–9 is linked to neurodegeneration in AD. Notably, female patients accounted for approximately half of the AD cohort, and the data from male and female patients were pooled together^89^. Here, we discovered that the levels of C7–C10 and undecanedioic (C11) acids were dramatically reduced in serum samples from patients with MCI, and even more reduced in females with MCI than in MCI males, with no differences between male and female controls. Several mechanisms could explain why we observed a profound decrease in the levels of dicarboxylic acids in MCI sera. For example, ∼60% of azelaic acid is eliminated unchanged in the urine. A portion undergoes β-oxidation to shorter chain dicarboxylic acids, and a part is decarboxylated^90^. These and other mechanisms could likely explain the lowered level of dicarboxylic acids in the serum of MCI patients. Still, it is unclear what mechanisms drive the sex-specific differences that we uncovered. In our study of metabolomic changes in severe COVID-19, suberic (C8) and sebacic (C10) acids were elevated in the serum of male and female patients, compared to non-infected controls, with no association of dicarboxylic acid levels with patient sex^91^. We conclude that dicarboxylic acid pathways behave in a disease-specific manner.

Phosphatidylcholines are phospholipids that contain choline. Previous studies reported phosphatidylcholine changes in AD. Three phosphatidylcholine species were lower in the plasma of patients with established AD than in cognitively healthy controls and MCI patients, and no sex association was observed^92^ (PC16:0, 20:5; PC16:0, 22:6; and 18:0, 22:6). A follow-up study in which ∼80% of AD patients were males also showed that specific phosphatidylcholine species were reduced in AD plasma^93^. The authors suggested that diminished phosphatidylcholine lipid levels could result from phosphatidylcholine hydrolysis induced by elevated phospholipase A2 activity^94^. Our experiments found 18 phosphatidylcholine species, many of which behave in unison and are higher in MCI male and female patients than control males and females. Most of these phosphatidylcholine lipids were higher in MCI female patients than MCI male patients.

Intriguingly, several saturated phosphatidylcholine lipid species positively correlate with tau phosphorylation in the CSF^95^. Phosphatidylcholine species again behaved in unison; the higher the levels of phosphorylated tau, the higher the levels of phosphatidylcholines, suggesting that saturated phosphatidylcholine lipids could be biomarkers of AD^95^. Phosphatidylcholines are a large and diverse family of lipids and could explain discrepancies between studies. Future studies should investigate how levels of specific phosphatidylcholines change at different stages of AD in both sexes.

Lysophospholipids modulate the immune system^96^. Therefore, these lipids may be related to the known differences in immune responses between males and females^97,98^. Our data show that levels of lysophospholipid metabolites were lower in male than female MCI patients, with few differences between sexes in the controls. Critically ill men and women exhibit profound disparities in metabolism^99^. In particular, the levels of lysophospholipids are higher in female patients than males^99^. We previously reported that lysophospholipids are reduced more in male patients with severe COVID-19 than in female patients, indicating that metabolites of lysophospholipids may be associated with sex in other age-dependent disorders^91^. Nevertheless, specific lysophospholipids may still be biomarkers for age-associated neurodegeneration since lysophospholipids are a large lipid family^100^, and various lysophospholipid metabolites and/or the enzymes that generate them vary between diseases^52^.

Long-chain fatty acids regulate a myriad of pathways in health and disease. The levels of many increase dramatically in metabolic disorders, such as diabetes, insulin resistance, and obesity^54^. In age-associated neurodegenerative disorders, disease-related changes in long-chain fatty acids are not fully understood. Several studies performed fatty acid blood lipidomics in MCI and AD^101–107^. A meta-analysis of these papers revealed no significant differences between controls and MCI patients for most fatty acids except docosahexaenoic acid (C22) and vaccenic acid (C18), which are, respectively, lower and higher in MCI individuals^108^. In AD, docosahexaenoic acid is lower than controls, but eicosatrienoic acid (C20) is elevated^108^. Overall, the data on fatty acids in MCI, and to a lesser extent in AD, may suffer from high variability due to various factors, including diet, often resulting in inconclusive outcomes. Our study detected 27 long-chain fatty acids in our samples, many of which are shifted in a specific direction in unison. For example, male MCI patients exhibit lower levels of many long-chain fatty acids compared to female MCI patients. Similarly, female MCI patients exhibit higher levels of many long-chain fatty acids compared to control female individuals. Taken together, long-chain fatty acid levels behave in unison in our data, suggesting that our observation of their changes is not random.

Glycerolipids are mono-, di-, and triesterified glycerols, corresponding to mono-, di-, and triacylglycerol, respectively. Monoacylglycerols are important metabolites and signaling molecules that regulate insulin secretion, food intake, autoimmune disorders, metabolic diseases, cancer, neurotransmission, and neurological diseases^55,56^. Triacylglycerols are not altered in the brain of AD patients, but both monoacylglycerols and diacylglycerols are elevated in the frontal cortex of MCI and AD patients^109–111^. Monoacylglycerols have sex-associated differences in metabolic diseases^112^, but their sex-dependent changes and roles in age-associated neurodegeneration remain poorly understood. Monoacylglycerols exist as 1-monoacylglycerols or 2-monoacylglycerols, depending on where the fatty acid is attached to the glycerol. 2 - monoacylglycerols have been extensively studied over the years, and some of 2-monoacylglycerols are endogenous cannabinoid receptor agonists^57^. 2-monoacylglycerols undergo isomerization to 1-monoacylglycerols, decreasing their cannabinergic potency. 1-monoacylglycerols are significantly less studied, but specific 1-monoacylglycerols regulate calcium signaling in thymocytes^113^, and 1-AG is an agonist of the TRPV1 channel^30^. We found that 1-palmitoleoyl glycerol and 1-AG were higher in female MCI subjects than in male MCI subjects with no differences between control males and females. In cultured male and female astrocytes, 1-AG promoted SphK2 phosphorylation, which was inhibited by TRPV1 receptor antagonists. We also showed that 1-AG promoted chromatin remodelling in cultured male and female astrocytes. The concentration differences of 1-monoacylglycerols between MCI males and females or sex-specific differences in TRPV1 channel signaling likely lead to sexually dimorphic mechanisms in MCI males and females.

In our study, we hypothesized that 2-monoacylglycerols were pro-senescence lipids that may affect cognition. We also hypothesized that 1-AG may activate SphK2, a lipid kinase that modulates transcription, amyloid formation, and cellular senescence^58,59^. Two sphingosine kinase enzyme isoforms, sphingosine kinase 1 (SphK1) and SphK2, synthesize sphingosine-1-phosphate by phosphorylating sphingosine^58^. SphK1 is a cytoplasmic enzyme; however, SphK2 can be localized to the nucleus, endoplasmic reticulum, or mitochondria. The activity of SphK2 is increased by phosphorylation^114^. In the nucleus, the SphK2/S1P pathway modulates transcription, epigenetic mechanisms, and telomere maintenance^58^. In particular, in cancerous cells, HDAC1 and HDAC2 associate with SphK2 in a transcription repressor complex^115^. S1P synthesized by SphK2 binds to and inhibits HDAC1 and HDAC2, thereby leading to an epigenetic regulation of transcription^115^. Increasing SphK2 levels in the neuronal nucleus leads to enhanced histone acetylation levels^116^. SphK2 is enriched in the neuronal nucleus in AD brains, compared to control samples^117^. In cystic fibrosis alveolar epithelial cells, SphK2 is hyperphosphorylated in the nucleus^114^. *Pseudomonas aeruginosa* infection in the lung epithelium results in SphK2 phosphorylation in the nucleus and enhanced acetylation of histones H3 and H4, which are important for the secretion of pro-inflammatory cytokines^114^. Our findings showed that, in cultured male and female astrocytes, 1-AG induced phosphorylation of SphK2 and more so in female cells. As a result, hyperactive SphK2 may trigger transcriptional changes and pro-senesence phenotypes in astrocytes in a sex-specific manner. This question is being pursued in our laboratory.

Chromatin covalent modifications, its organization, and architecture play a critical role in aging^118–120^. Condensed heterochromatic regions are reduced, and many nucleosomes are lost in aged cells. There are changes in DNA methylation and histone acetylation^118^. Proteins involved in genome 3D architecture, including nuclear laminins, have altered localization and expression in aged and diseased cells^119^. G-quadruplex DNA, a four-stranded higher-order DNA structure that folds from single-stranded guanine (G)-rich DNA sequences, has a role in aging^61^. G-quadruplexes are thermodynamically stable structures, with fast and reversible formation kinetics^121^, that are regulated in the cell by protein chaperones and helicases^122,123^. As a result, G-quadruplexes are ideal genetic modulators of DNA-dependent processes, including gene expression and replication^124,125^. G-quadruplex-forming motifs are often located in gene promoters^126^ and near the replication start sites^127^. Analyses of G-quadruplex DNA structures revealed that G-quadruplex DNA landscapes differ between cell types^128,129^. We previously demonstrated that neurons, astrocytes, and microglia basally exhibit different G-quadruplex landscapes^130^. We also showed that aged mouse brains contain more G-quadruplex DNA levels than young brains, and pharmacologically stabilizing G4-quadruplexes led to cognitive impairment in mice^131^, implicating these non-conventional DNA structures in aging^61^. This study showed that 1-AG modulates the G-quadruplex DNA landscape in astrocytes cultured from aged male and female mice in a sexually dimorphic fashion. Future studies will determine the molecular mechanisms that orchestrate lipid-dependent modulation of G-quadruplex DNA in healthy aging and in AD.

Our global metabolomics analyses demonstrated that MCI patients exhibited considerable metabolomic changes with several important sexually dimorphic pathways that could be developed as sex-specific biomarkers in AD if validated in larger cohorts. Phosphatidylcholine lipids, lysophospholipids, dicarboxylic fatty acids, long-chain fatty acids, and monoacylglycerol metabolites, as well as dipeptides and fibrinogen cleavage peptides and energy pathways, have an association with sex. These metabolites could serve as sex-specific biomarkers in MCI and/or AD and targeting these pathways could lead to important new therapeutic strategies.

## Declarations

### Ethics approval and consent to participate

Mice were maintained in accordance with guidelines and regulations of the University of Texas McGovern Medical School at Houston. All experimental protocols were approved by the University of Texas McGovern Medical School at Houston. The methods were carried out in accordance with the approved guidelines. Informed consent was obtained from all subjects.

### Consent for publication

Informed consent was obtained from all subjects.

### Availability of data and material

Metabolomics data are available in article supplementary material.

### Author Contributions

Data curation, R.D.E., M.J.V.K., G.J.O., A.M.G., H.F., P.H., M.P.B.C., G.D.C., H.W.A., L.C., S.H.K., M.H., C.M.F., B.P.G., P.E.S., L.D.M., A.S.T.; Supervision, S.P.M., A.U., B.P.G., P.E.S., L.D.M., A.S.T.; Writing – Original draft preparation, A.S.T.; Writing – Review & editing, R.D.E., M.J.V.K., A.M.G., G.J.O., S.H.K., M.H., C.F., B.P.G., S.P.M., A.U., P.E.S., L.D.M., A.S.T. All authors reviewed the manuscript.

## Acknowledgments

We thank Zheng Tan (Center for Healthy Aging, Changzhi Medical College, Changzhi, Shanxi, China) for allowing the use of the G4P protein^42^ in our studies. We thank members of the A.S.T. laboratory, the BRAINS laboratory for useful discussions, and the biorepository of neurological diseases team. Danielle Guillory provided administrative assistance. We thank a team of researchers from Metabolon for data generation, analyses, useful discussions, and administrative assistance: Scott McCulloch, Brian Ingram, Andrew Schwab, Madison Brooks, Gordon Irvine, and Justin Rogers.

## Supplemental figure and table legends

**Supplemental Table 1. Clinical characteristics of control individuals and MCI patients**. Serum samples from 20 male and 20 female control individuals and 20 male and 20 female MCI patients were used for analyses by ultrahigh-performance liquid chromatography-tandem mass spectroscopy.

**Supplemental Data 1**. The statistical heat map associated with the statistical analyses of the raw data. A biochemical name with no symbol indicates a compound confirmed based on an authentic chemical standard, and Metabolon is highly confident in its identity (Metabolomics Standards Initiative Tier 1 identification). A biochemical name with an * indicates a compound that has not been confirmed based on a standard, but Metabolon is confident in its identity. A biochemical name with an ** indicates a compound for which a standard is not available, but Metabolon is reasonably confident in its identity. A biochemical name with a (#) or [#] indicates a compound that is a structural isomer of another compound in the Metabolon spectral library. For example, a steroid that may be sulfated at one of several positions that are indistinguishable by the mass spectrometry data or a diacylglycerol for which more than one stereospecific molecule exists.

**Supplemental Data 2**. APOE genotypes of MCI individuals.

**Supplemental Data 3**. Association of metabolites with APOE genotypes and sex in the MCI cohort.

**Supplemental Data 4**. Association of metabolites with the MMSE scores in the MCI cohort.

**Supplemental Data 5**. Association of metabolites with MoCA scores in the MCI cohort.

**Supplemental Fig. 1. Global metabolomic profiling of male and female MCI patients and control individuals**

**(a)** PCA of metabolomic data for 20 male and female MCI patients and control subjects. Each dot represents an individual subject. The percentage values refer to the percentage of total variance associated with each component. Note how one individual separated from others. Purple: control male and female subjects, green: MCI male and female subjects. **(b)** PCA of metabolomic data for 20 male and female MCI patients and control subjects. Each dot represents an individual subject. The percentage values refer to the percentage of total variance associated with each component. Note how one individual separated from others. Blue: male control subjects and MCI patients, red: female control subjects and MCI patients.

**Supplemental Fig. 2. Volcano plots to illustrate metabolite differences**

Metabolites are plotted according to log2(fold-change) on the x-axis and -log10(P value) on the y-axis. Metabolites were considered significantly different if fold-changes (vertical dotted lines) were greater than 2 or less than -2 and false discovery rate corrected (FDR) p-values (horizonal dotted line) were less than 0.05. **(a)** The volcano plot illustrates metabolite differences between control females and control males. **(b)** The volcano plot illustrates metabolite differences between MCI female patients and MCI male patients.

**Supplemental Fig. 3. Succinate and phosphate exhibit an association with sex and MCI status**

Red and green cells indicate p≤0.05 (red indicates the fold-change values are significantly higher for that comparison; green values are significantly lower). Light red and light green shaded cells indicate 0.05<p<0.10 (light red indicates the fold-change values trend higher for that comparison; light green values trend lower). Note that there is a sex association for some metabolites.

**Supplemental Fig. 4. Purity of primary aged male and female astrocytes culture**

Astrocytes were fixed and stained with antibodies against AQP4 (red), S100β (green), and with the nuclear DAPI dye (blue). All cells were positive for AQP4 and S100β. Bar 10 µm.

**Supplemental Fig. 5. 1-AG enhances SphK2 phosphorylation in aged mouse male astrocytes**

**(a)** Male astrocytes were treated with control (vehicle), 1-AG alone (4 µM) or with TRPV1 inhibitors (2 μM CAY10568 and 2 μM JNJ17203212, overnight). Astrocytes were fixed and stained with antibodies against SphK2 (green), GFAP (red), and with the nuclear DAPI dye (blue), and imaged. Bar, 10 µm. **(b)** Male astrocytes were treated with control (vehicle), 1-AG alone (4 µM) or with TRPV1 inhibitors (2 μM CAY10568 and 2 μM JNJ17203212, overnight). Astrocytes were fixed and stained with antibodies against p-SphK2 (green), GFAP (red), and nuclear DAPI dye (blue), and imaged. Bar, 10 µm. **(c)** SphK2 intensities were measured from (a) with ImageJ (arbitrary units). **p=0.004, n.s., non-significant (one-way ANOVA). At least 250 cells were analyzed from three experiments. **(d)** p-SphK2 intensities were measured from (b) with ImageJ (arbitrary units). **p=0.005, ***p=0.0003, ****p<0.0001, n.s., non-significant (one-way ANOVA). At least 250 cells were analyzed from three experiments.

**Supplemental Fig. 6. 1-AG increases G-quadruplex-DNA levels in male and female astrocytes**

**(a)** Aged male and female astrocytes were infected with the G-quadruplex DNA reporter G4P-mScarlet. At 48 hours thereafter, astrocytes were treated with a vehicle or 1-AG (4 µM) overnight and imaged. Bar, 10 µm. On the left panel, zoomed images demonstrate G-quadruplex landscapes in individual cells. Bar, 30 µm. **(b)** The puncta index of the G4P-mScarlet reporter was measured in images from (a). *p =0.037, **p=0.0057, ****p<0.0001, n.s., non-significant (one-way ANOVA). More than 300 cells per condition were analyzed from four experiments.

**Supplemental Fig. 7. Differences in metabolism of sphingosines, sphingolipid synthesis, dihydroceramides, and ceramides**

Red and green cells indicate p≤0.05 (red indicates the fold-change values are significantly higher for that comparison; green values are significantly lower). Light red and light green shaded cells indicate 0.05<p<0.10 (light red indicates the fold-change values trend higher for that comparison; light green values trend lower). Note that there is a sex association for many metabolites.

**Supplemental Fig. 8. Differences in metabolism of sphingomyelins**

Red and green cells indicate p≤0.05 (red indicates the fold-change values are significantly higher for that comparison; green values are significantly lower). Light red and light green shaded cells indicate 0.05<p<0.10 (light red indicates the fold-change values trend higher for that comparison; light green values trend lower). Note that there is a sex association for many metabolites with no association with MCI status.

**Supplemental Fig. 9. Association of metabolites with the MMSE scores in MCI patients**

P values < 0.05 were counted as significant. Metabolites with a positive association (higher levels linked to higher MMSE scores) are indicated with red triangles. Those with a negative association (higher levels linked to lower MMSE scores) are indicated with blue triangles. Biochemical names followed by a number in parentheses indicate that a compound is a structural isomer of another compound in the spectral library.

**Supplemental Fig. 10. Association of metabolites with the MoCA scores in MCI patients**

P values < 0.05 were counted as significant. Metabolites with a positive association (higher levels linked to higher MoCA scores) are indicated with red triangles. Those with a negative association (higher levels linked to lower MoCA scores) are indicated with blue triangles. Biochemical names followed by a number in parentheses indicate that a compound is a structural isomer of another compound in the spectral library.

