## Supplementary figures and images for "Serum metabolome profiling in patients with mild cognitive impairment reveals sex differences in lipid metabolism"

### Supp Figure 1

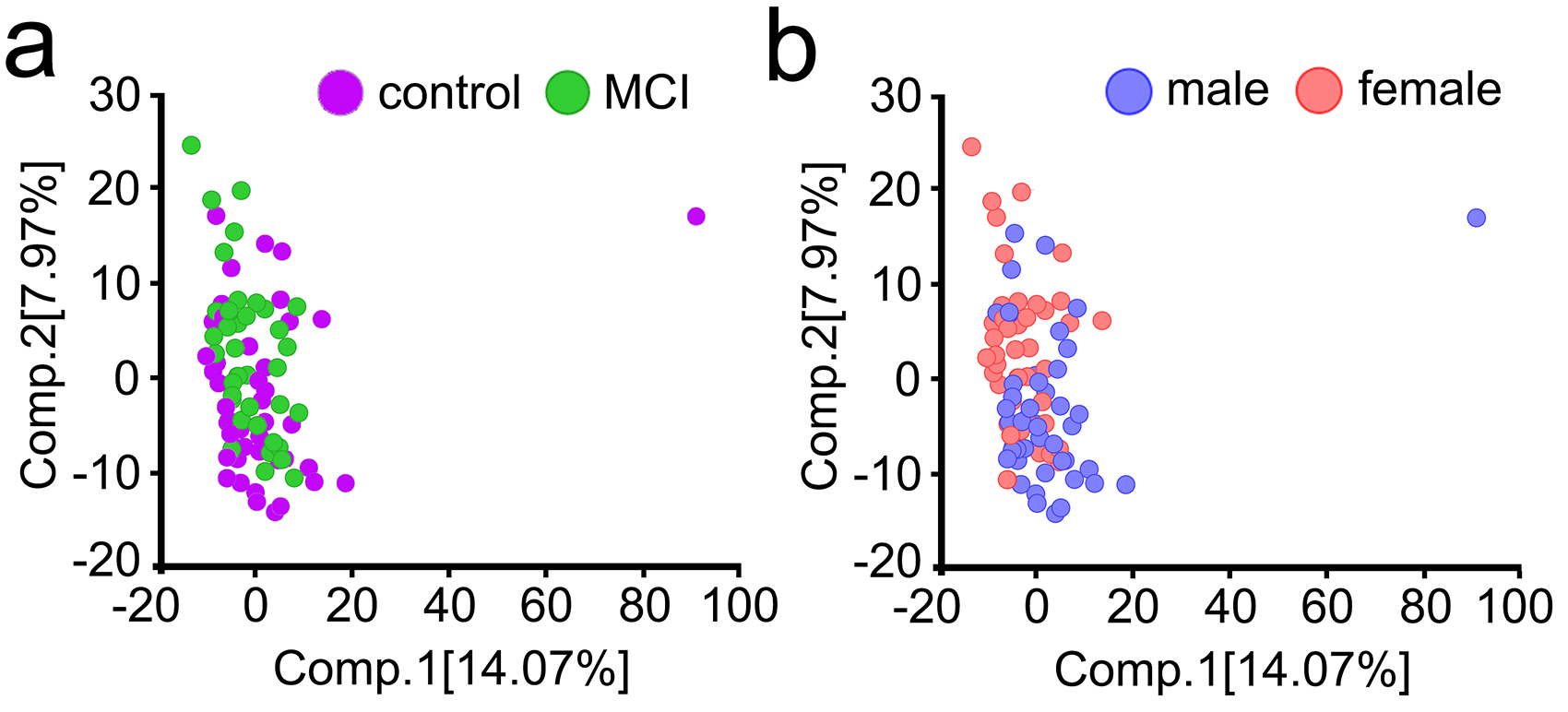

### Supp Figure 2

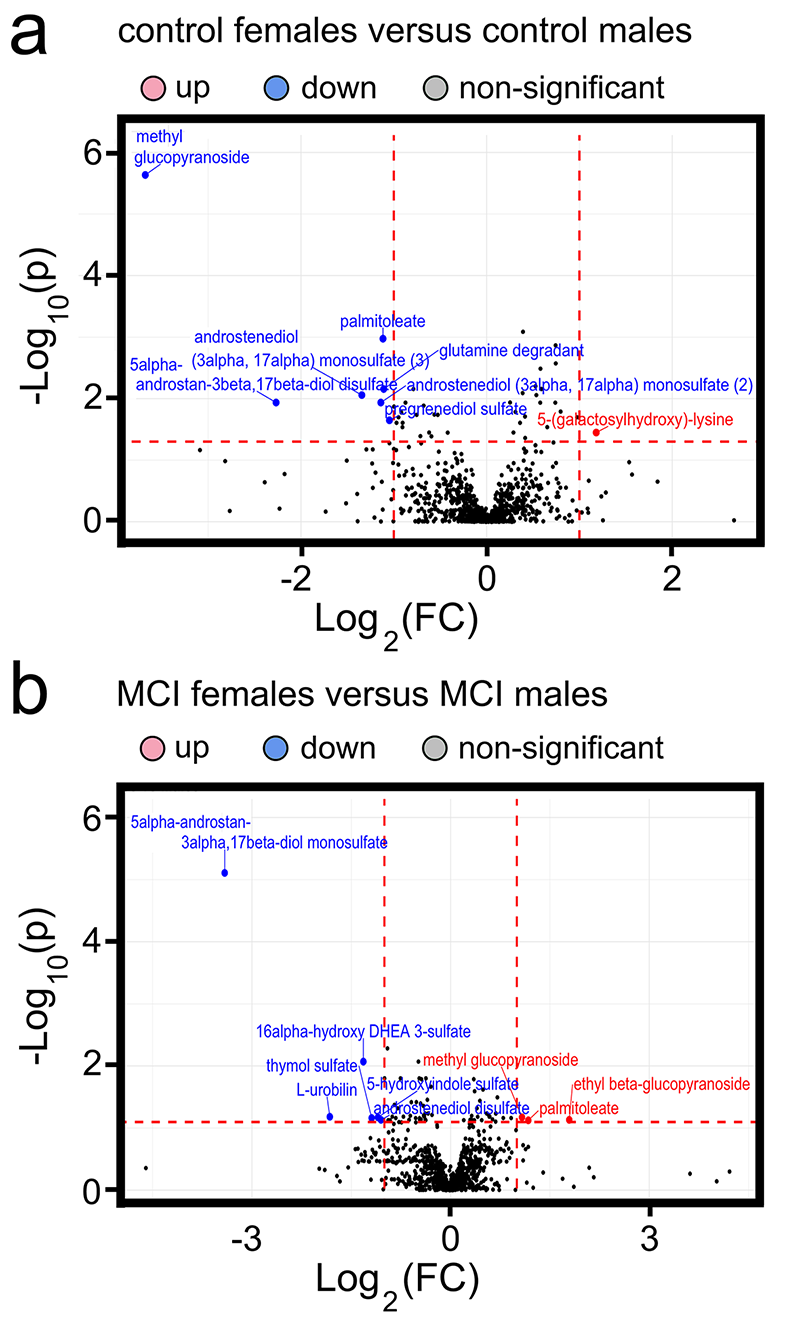

### Supp Figure 3

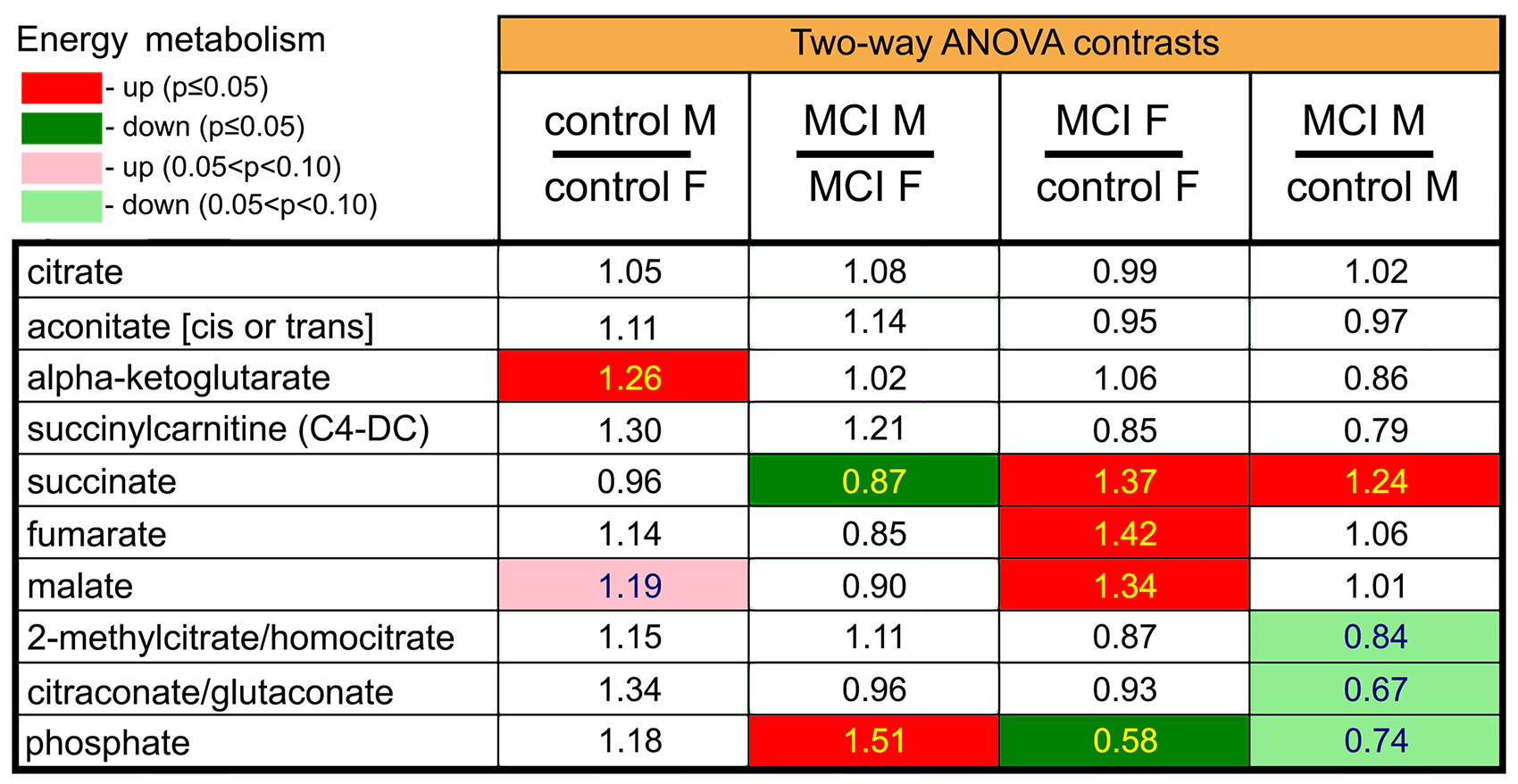

### Supp Figure 4

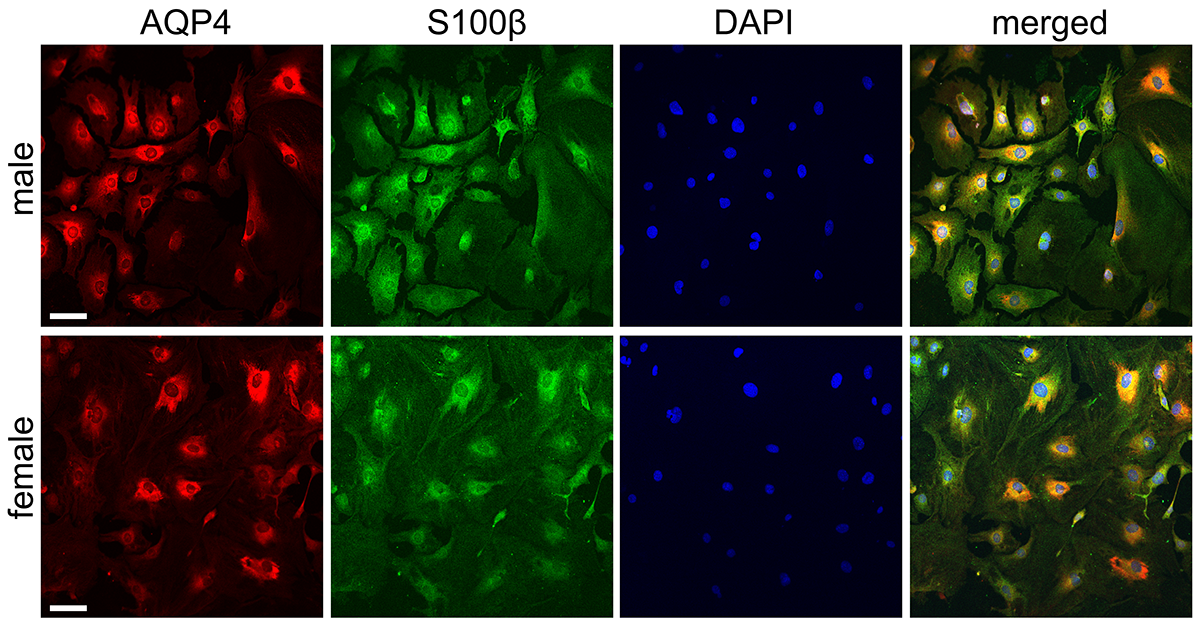

### Supp Figure 5

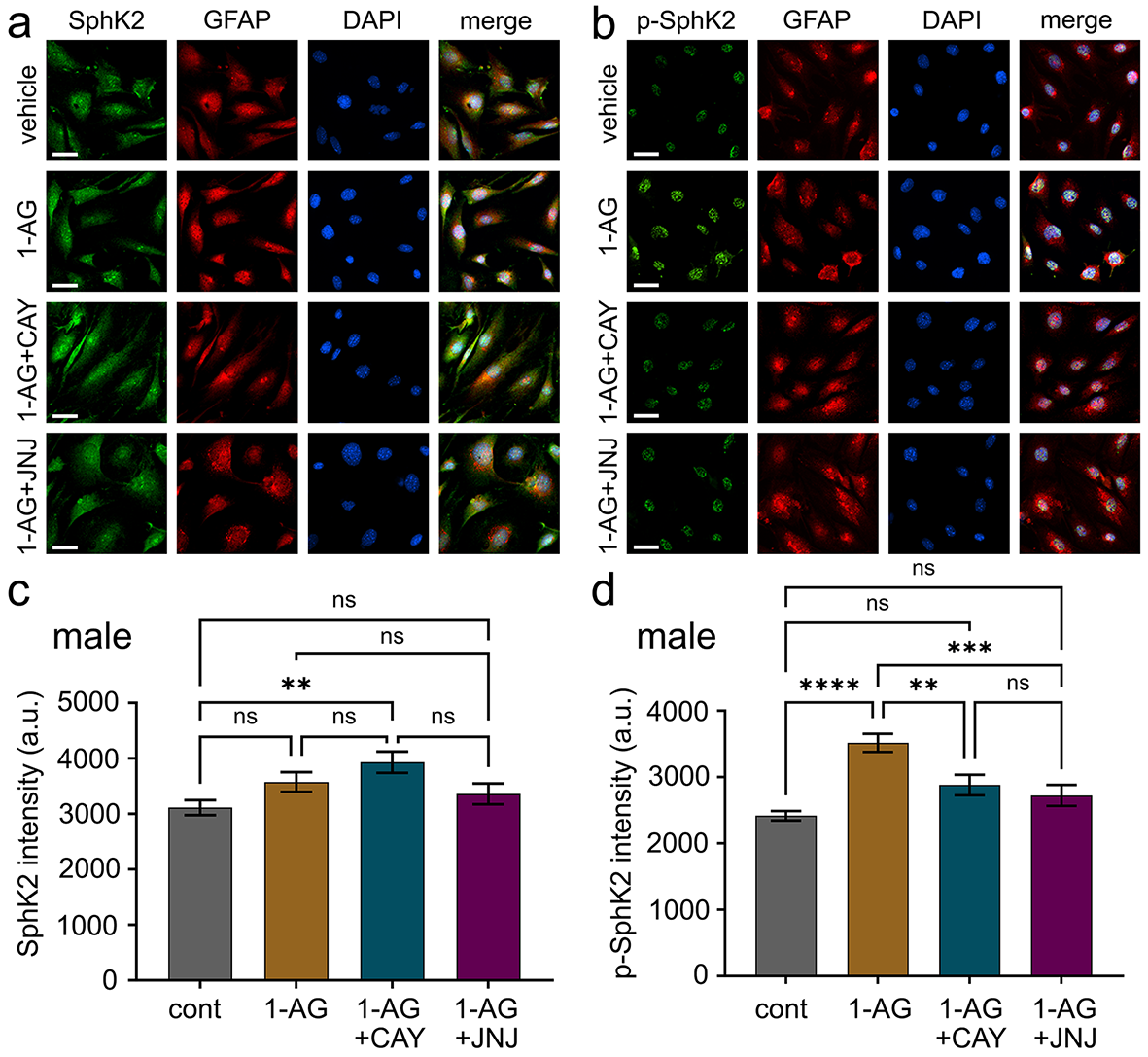

### Supp Figure 6

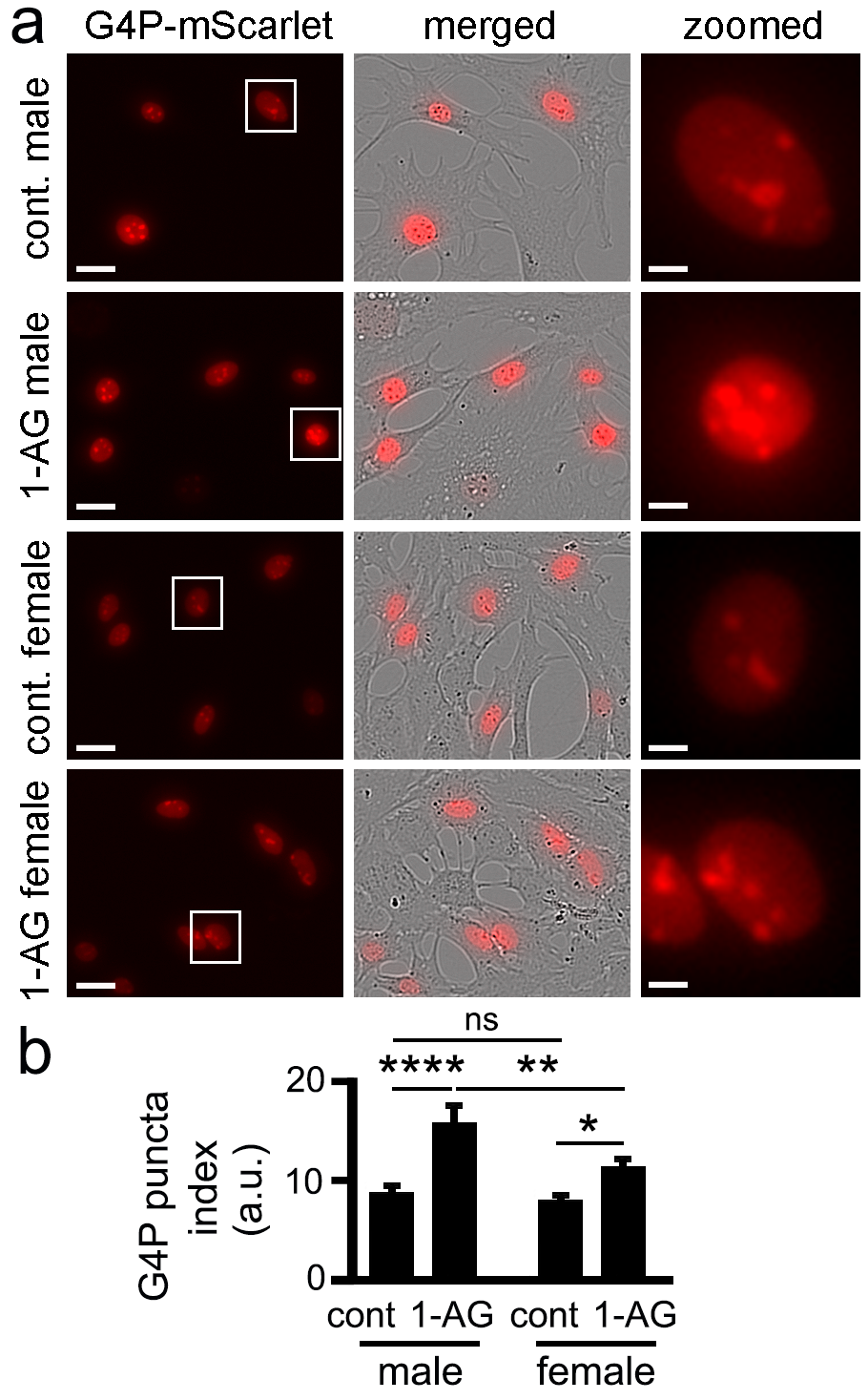

### Supp Figure 7

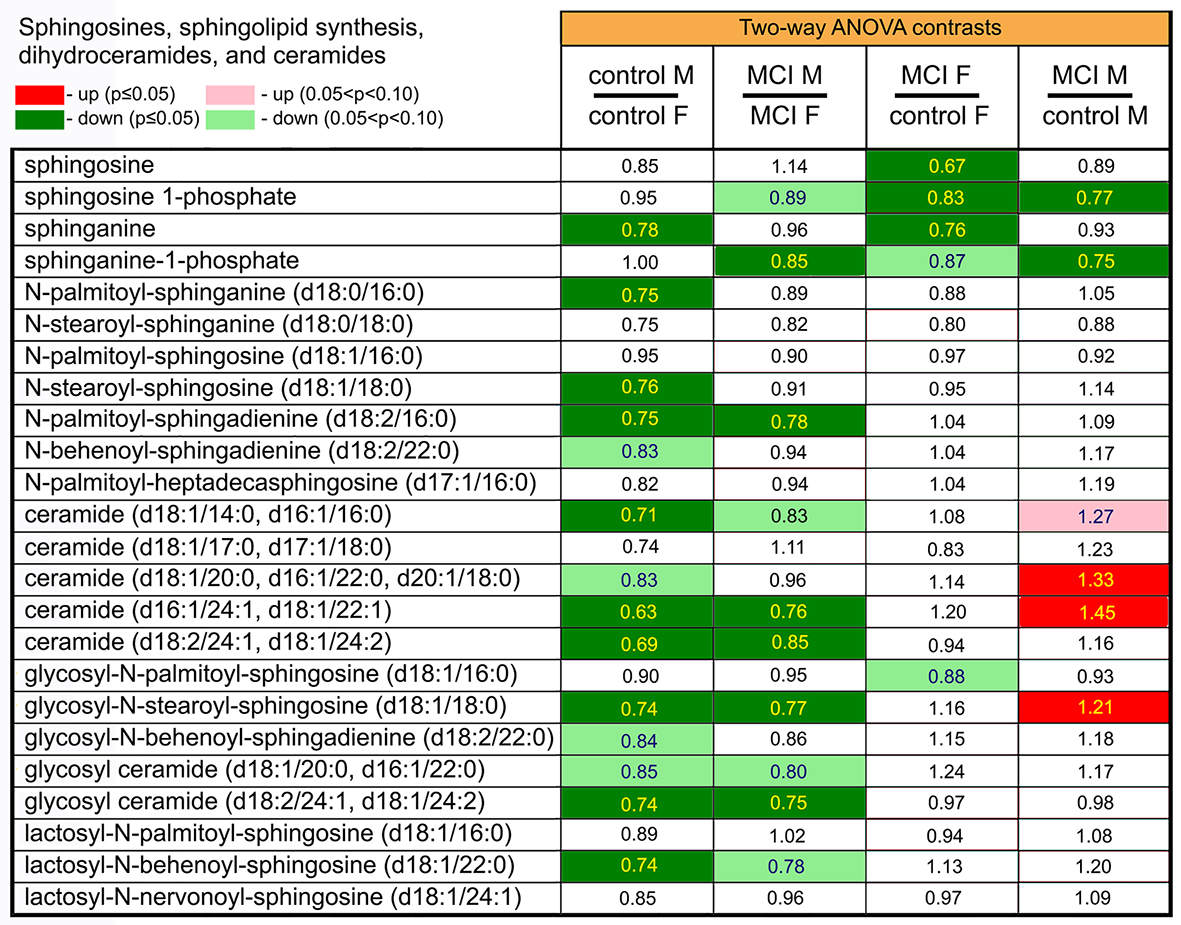

### Supp Figure 8

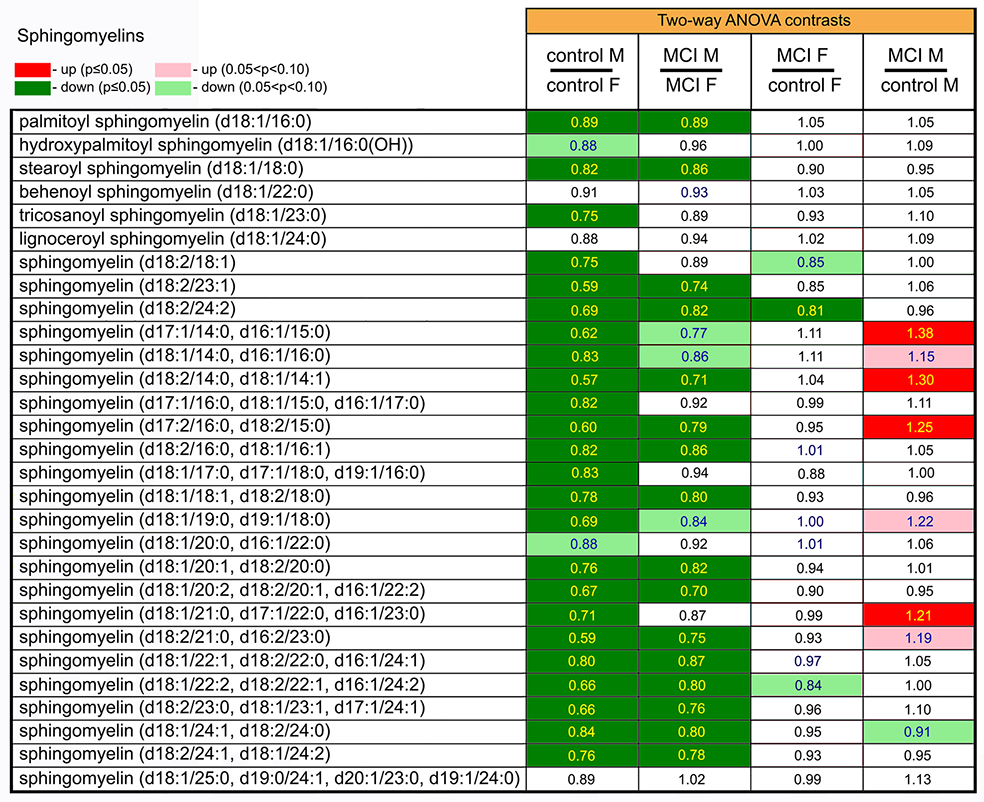

### Supp Figure 9

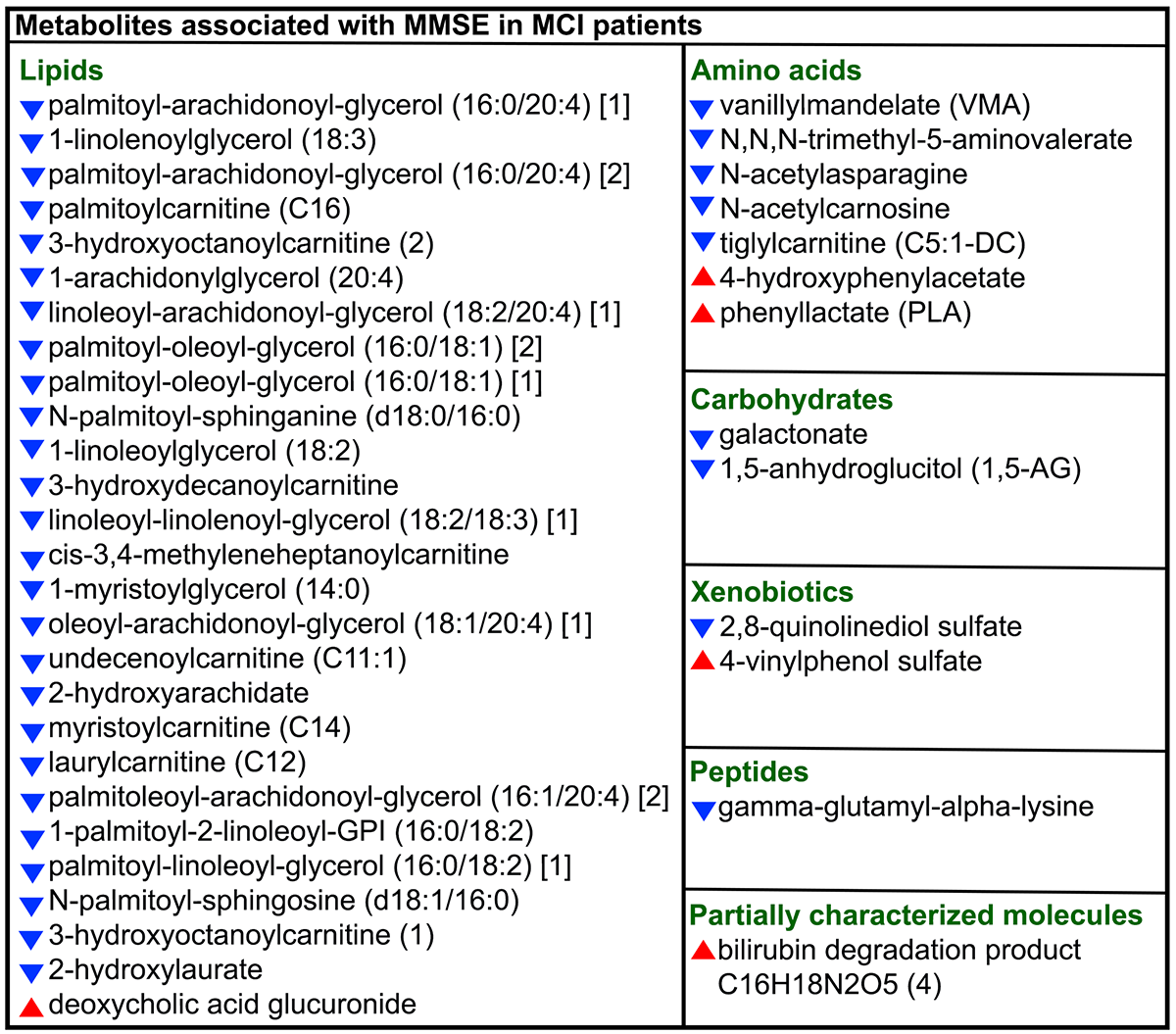

### Supp Figure 10

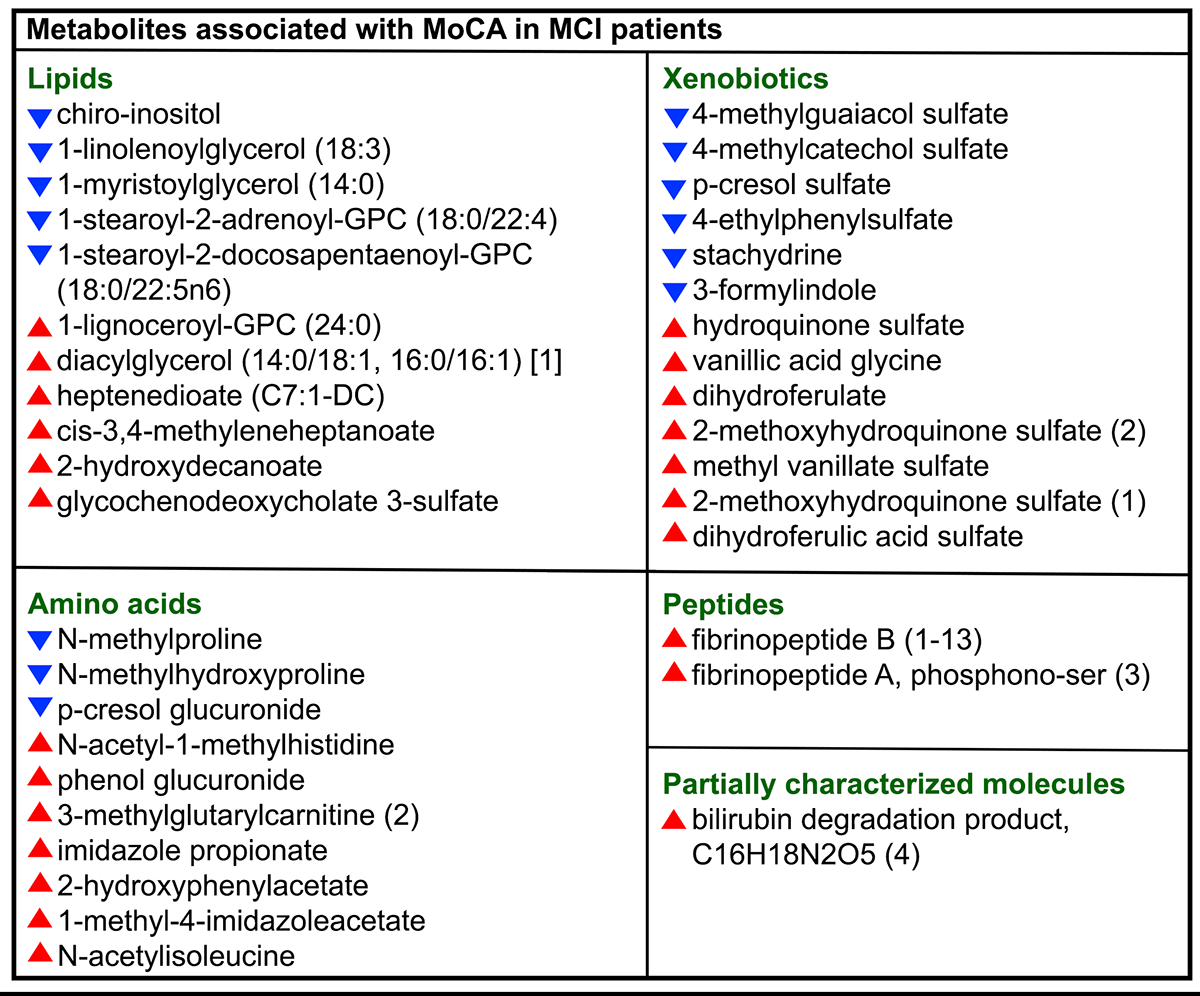
