## Supplementary material for "Serum metabolome profiling in patients with mild cognitive impairment reveals sex differences in lipid metabolism": Supp table 1

**Table 1. Clinical information of controls and MCI patients**

| Variable | Control (N=40) | | MCI (40) | |
|---|---|---|---|---|
| **Sex no. (%)** | | | | |
| Female | 20 (50.0) | | 20 (50.0) | |
| Male | 20 (50.0) | | 20 (50.0) | |
| **Age (year)** | Female | Male | Female | Male |
| Mean (SD) | 62.05 ± 12.98 | 61.95 ± 16.90 | 71.8 ± 5.75 | 74.2 ± 6.94 |
| Median (IQR) | 64 (72-50) | 63.5 (73.-47) | 72.5 (75-68) | 75.5 (78-73) |
| Range | 41-89 | 24-96 | 60-84 | 61-84 |
| **Race no. (%)** | | | | |
| Black | 6 (30.0) | 2 (10.0) | 1 (5.0) | 2 (10.0) |
| Hispanic | 1 (5.0) | 5 (25.0) | 1 (5.0) | 1 (5.0) |
| UTD | 0 (0.0) | 1 (5.0) | 0 (0.0) | 0 (0.0) |
| White | 13 (65.0) | 12 (60.0) | 18 (90.0) | 17 (85.0) |
| **BMI** | | | | |
| Mean (SD) | 28.81 ± 6.69 | 28.89 ± 5.32 | 24.37 ± 4.36 | 26.99 ± 4.24 |
| Median (IQR) | 30.54 (34.1-22.1) | 28.52 (29.2-27.0) | 23.6 (27-21) | 26 (29-23) |
| Range | 20.5-39.5 | 29.9-41.4 | 18.7-35.4 | 20.5-34.2 |
| **Comorbidity no. (%)** | | | | |
| Mean (SD) | 2.05 ± 1.70 | 2.35 ± 1.98 | 1.95 ± 1.35 | 2.70 ± 1.41 |
| Hyperlipidemia | 11(55.0) | 8 (40.0) | 16 (80.0) | 17 (85.0) |
| Hypertension | 10 (50.0) | 10 (50.0) | 9 (45.0) | 11 (55.0) |
| Obesity | 5 (25.0) | 2 (10.0) | 4 (20.0) | 5 (10.0) |
| Diabetes I | 2 (10.0) | 3 (15.0) | 0 (0.0) | 0 (0.0) |
| Diabetes II | 0 (0.0) | 2 (10.0) | 0 (0.0) | 2 (10.0) |
| History of cancer | 7 (35.0) | 6 (30.0) | 3 (15.0) | 4 (20.0) |
| Coronary Artery Disease | 1 (5.0) | 4 (20.0) | 0 (0.0) | 0 (0.0) |
| CHF | 2 (10.0) | 2 (10.0) | 0 (0.0) | 1 (5.0) |
| Carotid Stenosis | 1 (5.0) | 1 (5.0) | 0 (0.0) | 0 (0.0) |
| Atrial Fibrillation | 2 (10.0) | 2 (10.0) | 0 (0.0) | 1 (5.0) |
| Ischemic Stroke | 0 (0.0) | 1 (5.0) | 0 (0.0) | 0 (0.0) |
| Intracerebral Hemorrhage | 0 (0.0) | 1 (5.0) | 0 (0.0) | 0 (0.0) |
| Stroke (type unknown) | 0 (0.0) | 2 (10.0) | 0 (0.0) | 0 (0.0) |
| ESRD | 0 (0.0) | 1 (5.0) | 0 (0.0) | 0 (0.0) |
| TBI | 0 (0.0) | (0.0) | 2 (4.0) | 6 (11.0) |
| Smoker | 0 (0.0) | 2 (10.0) | 5 (25.0) | 7 (35.0) |

no. (%), number; SD, standard deviation; IQR, interquartile range; Congestive Heart Failure, CHF; End Stage Renal Disease, ESRD; UTD, unable to determine.
